# Limited Environmental Serine Confers Sensitivity to PHGDH Inhibition in Brain Metastasis

**DOI:** 10.1101/2020.03.03.974980

**Authors:** Bryan Ngo, Eugenie Kim, Victoria Osorio-Vasquez, Sophia Doll, Sophia Bustraan, Alba Luengo, Shawn M. Davidson, Ahmed Ali, Gino D. Ferraro, Diane Kang, Jing Ni, Roger Liang, Ariana Plasger, Edward R. Kastenhuber, Roozbeh Eskandari, Sarah Bacha, Roshan K. Siriam, Samuel F. Bakhoum, Edouard Mullarky, Matija Snuderl, Paolo Cotzia, Nello Mainolfi, Vipin Suri, Adam Friedman, Mark Manfredi, David M. Sabatini, Drew Jones, Min Yu, Jean J. Zhao, Rakesh K. Jain, Matthew G. Vander Heiden, Eva Hernando, Matthias Mann, Lewis C. Cantley, Michael E. Pacold

**Author notes:** These Authors Contributed Equally (Bryan Ngo, Eugenie Kim, Victoria Osorio-Vasquez). Correspondence and requests for materials should be addressed to: M.E.P. and/or L.C.C.

## Abstract

A hallmark of metastasis is the adaptation of tumor cells to new environments. Although it is well established that the metabolic milieu of the brain is severely deprived of nutrients, particularly the amino acids serine and its catabolite glycine, how brain metastases rewire their metabolism to survive in the nutrient-limited environment of the brain is poorly understood. Here we demonstrate that cell-intrinsic *de novo* serine synthesis is a major determinant of brain metastasis. Whole proteome comparison of triple-negative breast cancer (TNBC) cells that differ in their capacity to colonize the brain reveals that 3-phosphoglycerate dehydrogenase (PHGDH), which catalyzes the rate-limiting step of glucose-derived serine synthesis, is the most significantly upregulated protein in cells that efficiently metastasize to the brain. Genetic silencing or pharmacological inhibition of PHGDH attenuated brain metastasis and improved overall survival in mice, whereas expression of catalytically active PHGDH in a non-brain trophic cell line promoted brain metastasis. Collectively, these findings indicate that nutrient availability determines serine synthesis pathway dependence in brain metastasis, and suggest that PHGDH inhibitors may be useful in the treatment of patients with cancers that have spread to the brain.

**Statement of Significance:** Our study highlights how limited serine and glycine availability within the brain microenvironment potentiates tumor cell sensitivity to serine synthesis inhibition. This finding underscores the importance of studying cancer metabolism in physiologically-relevant contexts, and provides a rationale for using PHGDH inhibitors to treat brain metastasis.

## Introduction

Metastasis is an evolutionary process whereby the adaptive potential of tumor cells enables their ability to colonize specific tissues.^1^ Despite efforts to identify novel oncogenic mutations that drive metastasis, whole-genome sequencing studies have revealed striking similarities in the array of driver gene mutations between primary tumors and matched metastases.^2,3^ These findings support the proposition that metastatic competence can arise without additional driver mutations and that metabolic adaptation, dictated by the selective pressure of the metastatic niche, may have a more profound role in the colonization of distant organs and response to therapy. Thus, investigating the dynamic interplay between environmental nutrient availability and cancer cell metabolism offers an attractive opportunity to identify and therapeutically target metabolic pathways that promote metastatic colonization of specific tissues, such as the brain.

Brain metastasis (BrM) is the most prevalent intracranial malignancy in adults.^4,5^ Current therapies for brain metastasis include surgical excision, stereotactic radiosurgery, whole brain radiotherapy (WBRT), and corticosteroids for symptom management. Yet, despite advancements in radiosurgery and immunotherapy, the median survival for patients with brain metastases rarely exceeds 12 months.^6-8^ An improved understanding of the characteristics and pathogenesis of brain metastasis would facilitate the development of more effective therapies to treat and manage cancers that have spread to the brain.

Growing evidence indicates that brain metastases possess unique molecular and metabolic characteristics compared with metastases at other sites.^4,9-15^ A classical view argues that the establishment and growth of brain metastases is in part dependent on cooperative cellular interactions between tumor cells and stromal cells (astrocytes, microglial and endothelial cells), within the metastatic niche.^9,10,14,16,17^ However, while much has been learned about the cellular and molecular determinants of brain metastasis, little is known about how cancer cells adapt to the compositionally simple nutrient microenvironment of the brain, which is nearly devoid of proteins and most amino acids.

Nutrient availability has emerged as an important regulator of cancer cell metabolism.^18-29^ In the brain, the interstitial fluid (ISF) occupies 25-30% of the total brain volume, and plays a crucial role in nutrient transport, waste product removal, and signal transmission among various cell types, including neurons, astrocytes, pericytes, and endothelial cells. With the exception of glutamine, both the brain ISF and CSF are severely depleted in proteins and amino acids relative to plasma, and are effectively comparable in composition.^30^ Despite the importance of nutrient availability in regulating cancer cell metabolism, it is unknown whether differences in interstitial brain and plasma nutrient concentrations enforce specific metabolic dependencies that alter cellular responses to metabolically targeted therapies.

Metabolic dependencies on specific amino acids vary remarkably with tissue of origin, oncogenic mutations, and environmental nutrient availability.^19,23,29^ Amino acids contribute to a diverse array of processes, and are important for cell proliferation and the biosynthesis of proteins, nucleotides, lipids, glutathione, and tricarboxylic acid (TCA) cycle intermediates. When amino acids are limiting, cells modulate the expression of specific transporters, or take up nutrients in bulk from the extracellular space through macropinocytosis.^22,31^ In addition, cells can synthesize a variety of non-essential amino acids (NEAAs), including serine, glycine, and aspartate, through *de novo* biosynthetic pathways. For example, diversion of glycolytic intermediates into the serine and glycine biosynthesis pathway is commonly observed in melanoma and triple-negative breast cancer (TNBC; ER^-^/PR^-^/Her2^-^).^32,33^ This specific metabolic alteration is driven by the increased expression of 3-phosphoglycerate dehydrogenase (PHGDH) through focal genomic amplifications, transcriptional changes, or post-translational regulation.^32-35^ However, attempts to therapeutically target PHGDH with small-molecules have largely yielded low potency inhibitors with modest effects *in vivo*.^36-39^ This raises the possibility that while cancer cell-intrinsic factors, such as genetic mutations, have the potential to dictate metabolic dependencies, the phenotypic penetrance of these events are determined by the availability of specific nutrients in the environment.

Here we investigate the mechanisms contributing to metastatic cancer cell survival within the brain microenvironment, and propose that amino acid availability has a profound influence on metabolic dependency and drug response in brain metastases. Using tailored cell culture media that reflect brain interstitial amino acid concentrations and refined mouse models of brain metastasis, we demonstrate that PHGDH is a major determinant and metabolic liability in brain metastasis. PHGDH activity is conditionally essential for cell proliferation in serine- and glycine-limited tissue environments. These environmental conditions enforce tumor dependency on *de novo* serine synthesis, and can be exploited by genetic or pharmacological inhibition of PHGDH. The selectivity of these results indicates that identical cells growing in different environments exhibit distinct metabolic programs and variation in drug sensitivity, and suggest that improved PHGDH inhibitors could be effective in treating patients with brain metastases.

## Results

### Generation and Characterization of Aggressive and Indolent Brain Metastatic Cells

Generation of brain-specific metastatic clones that differ in their ability to colonize the brain through the *in vivo* selection of cancer cell lines permits the direct experimental interrogation of traits that promote brain metastasis.^9,10,14,17,40^ As a source, we selected the TNBC cell line, MDA-MB-231, because of its proclivity to engraft and metastasize after hematogenous dissemination induced by intracardiac administration (Fig. 1A). Cells were engineered to stably express luciferase and GFP, inoculated directly into the left ventricle of immunodeficient athymic nude mice, and monitored weekly by bioluminescence imaging (BLI). Brain metastatic cells were isolated by fluorescence activated cell sorting (FACS) and recovered in culture. Recovered cells were reinjected into the arterial circulation of mice, and this process was repeated three times to generate aggressive and indolent cell lines that possessed the ability to reach the brain from the circulation. Indolent brain metastatic cells were isolated from mechanically dissociated BLI-negative brains. Unlike highly aggressive brain metastatic clones, which generate multifocal lesions in the cerebrum, cerebellum, and brain stem, mice injected with indolent brain metastatic cancer cells remain healthy for ∼10 weeks (2.5 months) without any detectable signs of brain metastasis progression by BLI (Fig. 1B, C; Supplementary Fig. S1A).

**Figure 1:**
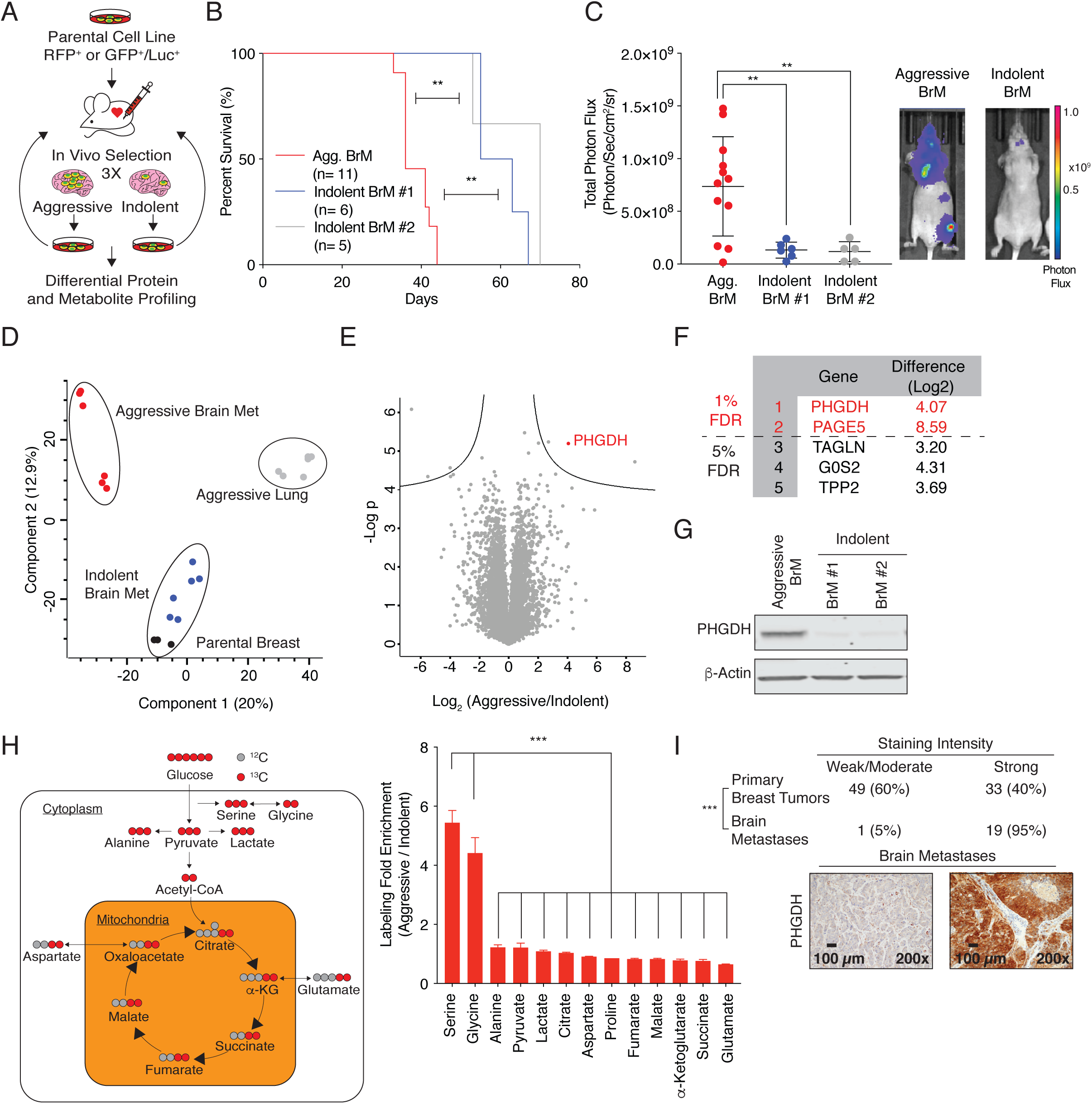
Generation and Characterization of Aggressive and Indolent Brain Metastatic Cells. **(A)** Schematic of the selection of brain metastatic (BrM) cell lines. Brain-trophic cells were selected through three rounds of intracardiac injection followed by sorting of cells isolated from the brains of mice. **(B)** Kaplan-Meier plot showing disease-specific survival of mice injected with aggressive (black) or indolent (blue and grey) brain metastatic cells. Significance was tested using the log-rank test. Mice injected with indolent brain metastatic cell lines live on average two weeks longer than mice injected with aggressive brain-trophic cells. **(C)** Average whole animal bioluminescence imaging (BLI) (photons/second) four weeks after intracardiac injection of aggressive and indolent brain metastatic cells. Aggressive brain-trophic cells have a higher brain burden than indolent cells. **(D)** Principal component analysis (PCA) plot of proteome derived from two aggressive brain metastatic clones (black), two aggressive lung metastatic cells (purple and light blue), two indolent brain metastatic cells (blue and grey), and parental MDA-MB-231 TNBC cells (green). The brain-trophic cells are distinct from the parental, indolent, and lung-trophic cells. **(E)** Volcano plot of the p-value vs. the log_2_ protein abundance differences between aggressive and indolent brain metastatic cells. Solid lines indicate FDR < 0.01. **(F)** List of the five most differentially expressed proteins in aggressive brain metastatic cells relative to indolent brain metastatic cells. PHGDH is the most differentially expressed gene with the lowest FDR. **(G)** Immunoblot of PHGDH expression in aggressive and indolent brain metastatic cells. PHGDH is more strongly expressed in aggressive cells than in the indolent cells. **(H)** Fold enrichment of the fractional labeling of metabolites by U-^13^C-glucose in aggressive vs. indolent brain metastatic cells. Glycine and serine have the highest fold enrichment of glucose-derived carbon of the examined metabolites in aggressive brain-trophic cells in comparison to indolent cells. **(I)** Representative immunohistochemical staining of PHGDH expression in human breast cancer brain metastases and primary breast tumors (Scale bar, 200 *μ*m). PHGDH is strongly expressed in human breast cancer brain metastases in comparison to a previously published cohort of primary breast cancers. Significance was determined using Fisher’s exact test. For this and all subsequent figures, * indicates statistical significance with p<0.05, ** indicates p<0.005, and *** indicates p<0.0005. Significance is measured with the Mann-Whitney *U* test unless otherwise indicated.

### Serine Synthesis is Upregulated in Aggressive Brain Metastatic Cells

To gain insight into the molecular events underlying brain metastasis, we analyzed both the peptide mixtures and labeling of central carbon metabolites from parental, aggressive, and indolent TNBC brain metastatic cells.^41-45^ Comparative proteomic expression profiling of TNBC cells that differ in their capacity to colonize the brain or lung using an optimized, high-sensitivity, label-free protein expression profiling workflow quantified ∼7,000 proteins per sample with excellent reproducibility.^44-47^ Our analysis yielded protein expression profiles of the parental, indolent, and aggressive brain metastatic cells that segregated independently by principal component analysis, indicating that each population is phenotypically distinct (Fig. 1D). 3-phosphoglycerate dehydrogenase (PHGDH), the first enzyme in the glucose-derived serine biosynthesis pathway, was the most significantly upregulated protein in aggressive brain metastatic cells compared to the non-aggressive brain metastatic cells (FDR < 0.01; Fig. 1E-G; Supplementary Fig. S1B, C). In parallel, we also labeled aggressive and indolent brain-trophic cells with glucose labeled at all six carbons with ^13^C (U-^13^C-glucose) and measured the enrichment of glucose-derived metabolites by gas chromatography-mass spectrometry (GC-MS). Interestingly, glucose-derived serine and glycine exhibited the greatest increase in glucose-derived ^13^C in aggressive brain metastatic cells compared to indolent brain metastatic cells (Fig. 1H). Taken together, our untargeted proteomics and U-^13^C-glucose tracing results both indicate that glucose-derived serine synthesis is greatly enhanced in cells that readily form brain metastasis.

### PHGDH is Enriched in Human Brain Metastases

PHGDH catalyzes the first and rate-limiting step in the canonical, glucose-derived serine synthesis pathway, and uses NAD+ as a cofactor to oxidize the glycolytic intermediate 3-phosphoglycerate to 3-phosphohydroxypyruvate (3-PHP) (Fig. 2A). Subsequent enzymes in the pathway convert 3-PHP into serine via transamination (phosphoserine aminotransferase 1, PSAT1) and phosphate ester hydrolysis (phosphoserine phosphatase, PSPH). Serine is essential for synthesis of proteins and other biomolecules needed for cell proliferation, including glycine, cysteine, nucleotides, phosphatidylserine, sphingosine, and charged folates required for one-carbon metabolism.^48^

**Figure 2:**
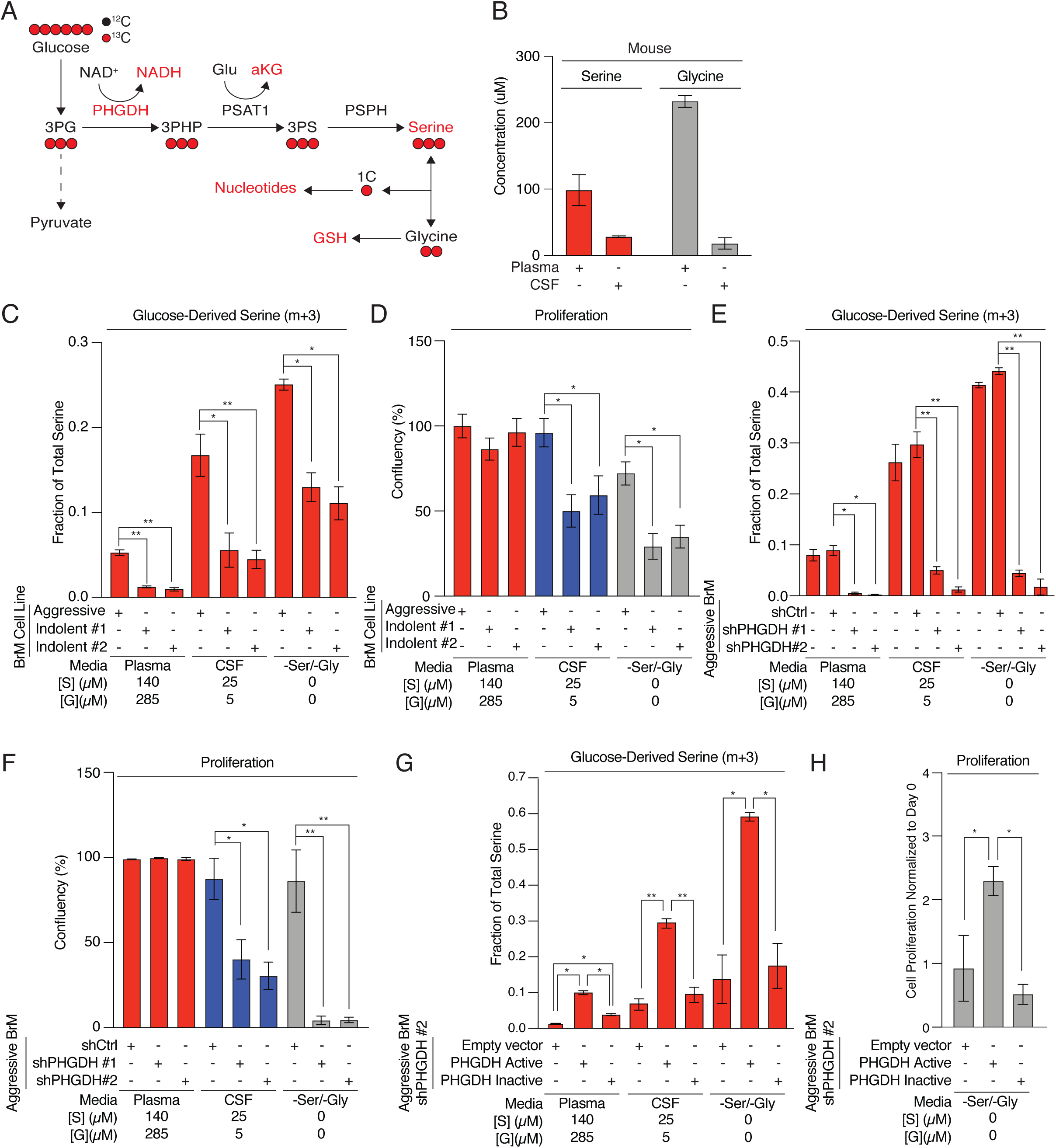
Aggressive Brain Metastases are Serine Prototrophs. **(A)** Schematic of the glucose-derived serine biosynthesis pathway. 3PHP (3-phosphohydroxypyruvate); 3PS (3-phospho-serine); 1C (one-carbon unit); GSH (reduced glutathione); PHGDH (phosphoglycerate dehydrogenase); PSAT (phospho-serine aminotransferase); PSPH (phosphoserine phosphatase). Glucose-derived carbons are highlighted in red. **(B)** Concentrations of serine and glycine in plasma or CSF collected from 6-week old female mice. **(C)** Fractional labeling of intracellular glucose-derived m+3 serine from aggressive or indolent brain metastatic cells grown Plasma, CSF, or -Ser/-Gly media. Aggressive brain metastatic cells synthesize more glucose-derived serine than indolent brain metastatic cells as indicated by the higher fraction of m+3 labeled serine. **(D)** Proliferative capacity of aggressive or indolent brain metastatic cells grown in Plasma, CSF, or -Ser/-Gly media. Aggressive and indolent brain metastatic cells proliferate identically in media with plasma concentrations of serine and glycine, but indolent brain metastatic cells do not proliferate as well as aggressive brain metastatic cells in CSF or -Ser/-Gly media. **(E)** Fractional labeling of intracellular glucose-derived serine from aggressive brain-trophic cell lines expressing a control shRNA (shCtrl) or PHGDH shRNA (shPHGDH #1 or shPHGDH #2). Cells with suppressed PHGDH synthesize less glucose-derived serine than wild-type cells or cells transduced with a control hairpin. **(F)** Proliferative capacity of aggressive brain metastatic cells expressing a control shRNA (shCtrl) or PHGDH shRNA (shPHGDH #1 or shPHGDH #2). PHGDH knockdown cells grow more slowly in CSF and -Ser/-Gly media than cells expressing a control shRNA. **(G)** Fractional labeling of intracellular glucose-derived serine from aggressive brain metastatic cells expressing either a catalytically active or catalytically inactive, shRNA-resistant PHGDH. Genetic rescue with catalytically active, shRNA-resistant PHGDH restores glucose-derived serine synthesis, while expression of catalytically inactive PHGDH fails to completely restore serine synthesis. **(H)** Proliferative capacity of PHGDH knockdown aggressive brain metastatic cells expressing either a catalytically active or catalytically inactive, shRNA-resistant PHGDH. Error bars represent standard deviations. Cell proliferation data was monitored over 4-6 days using an Incucyte or Multisizer Coulter Counter.

Previous studies have shown that PHGDH is selectively amplified in human TNBC and melanoma, and is associated with poor patient survival.^32,33,49,50^ To reveal if PHGDH expression is enhanced in human brain metastases, we determined PHGDH expression levels by immunohistochemistry from tumor biopsies of breast cancer brain metastases (n=20), and compared these to primary breast tumor biopsies.^32^ Nineteen out of twenty patients (95%) had brain metastases that strongly expressed PHGDH (Fig. 1I). This is a significantly higher percentage than the ∼40% of patients with primary breast tumors that strongly expressed PHGDH as reanalyzed from a previously published cohort.^32^ In addition, isolation of patient derived estrogen-positive (ER^+^) circulating tumor cells (CTCs) that exhibited tropism towards the brain (BrM3) had a dramatic increase in PHGDH expression compared to initial CTCs injected into mice (Brx50 Parental) (Supplementary Fig. S1D, E). Analysis of Oncomine data also indicated that PHGDH expression is significantly higher in TNBC tumors (n=46) than in other breast cancer subtypes (n =250) (from TCGA), and PHGDH expression in breast cancer with metastatic events at 3 years (n = 140) is higher than breast cancer without metastatic events at 3 years (n = 48) (Supplementary Fig. S1f, g).^10^ Taken together, these data suggest that a higher fraction of brain metastases express PHGDH than primary tumors, and that the poorer outcomes of patients with PHGDH-overexpressing tumors may be due to higher risk of metastasis, including brain metastasis.^32,33^

### The Brain is a Serine- and Glycine-Limited Environment

Published data indicate that the concentrations of most proteogenic amino acids in the brain ISF and CSF are comparable, and are substantially lower than in the plasma (Supplementary Fig. S2A).^30,51^ For this reason, we use the amino acid concentrations found in CSF as a proxy for brain ISF, and postulated that upon vascular extravasation, cancer cells that colonize the brain likely encounter an environment with low concentrations of amino acids. To confirm that amino acid concentrations differ between the plasma and CSF in mice, we quantified proteogenic amino acids from paired mouse CSF and plasma samples using GC-MS. In agreement with previous studies, the concentration of serine and glycine in mouse CSF is ∼20uM and ∼5uM, respectively, about 5- and 50-fold lower than in mouse plasma (100 and 246 μM, respectively) (Fig. 2B).^30^

This observation, in combination with our proteomic and metabolomic data showing enrichment of serine synthesis in highly aggressive brain metastatic cells, prompted us to ask whether the nutrient-limited environment of the brain might select for cancer cells that are serine prototrophs. As both serine and glycine are proteinogenic amino acids and are the source of one-carbon units that are necessary for cell growth, but are among the least abundant amino acids in the CSF, we hypothesized that serine prototrophy enables tumor cell colonization of the brain.

### PHGDH is Required for Serine Synthesis and Cell Proliferation in Serine- and Glycine-Limited Environments

To understand the consequences of limited serine and glycine availability on cancer cell metabolism and proliferation in the brain metastatic setting, we developed cell culture medium with serine and glycine concentrations comparable to plasma and CSF (Plasma, CSF, and -Ser/-Gly media) (Supplementary Fig. S2B). We then measured the incorporation of U-^13^C-glucose into serine and glycine in aggressive and indolent brain metastatic cells cultured in plasma, CSF, or - Ser/-Gly media. The fraction of glucose-derived serine (m+3) was higher in both aggressive and indolent brain metastatic cells cultured in CSF media and -Ser/-Gly media (Fig. 2C; Supplementary Fig. S2C, D). PHGDH-expressing aggressive brain metastatic cells derived 5% of their serine from glucose. This increased to ∼20% in CSF media and ∼25% in -Ser/-Gly media. In contrast, non-aggressive brain-trophic cell lines derived no more than ∼15% of their serine from glucose regardless of the extracellular serine and glycine concentrations (Fig. 2C).

As serine is essential for the production of one-carbon units needed cell proliferation, and can be taken up from the extracellular environment through amino acid transporters, we predicted both aggressive and indolent brain metastatic cell lines should proliferate at comparable rates in plasma media. However, only PHGDH-expressing, aggressive brain metastatic cells that are capable of synthesizing serine from glucose, would proliferate in serine- and glycine-limited conditions (CSF and -Ser/-Gly media). Measurements of cell proliferation using an automated live cell analysis platform (Incucyte, Essen BioScience) confirmed that aggressive brain metastatic cells, expressing PHGDH, proliferated in CSF and -Ser/-Gly media, whereas indolent brain metastatic cancer cells with decreased baseline PHGDH expression, proliferated poorly when cultured in these media conditions (Fig. 2D). Thus, cells with elevated PHGDH and high serine synthesis pathway flux were capable of robust proliferation in media lacking serine and glycine, whereas cells with low PHGDH expression proliferate poorly in serine- and glycine-low conditions typically found in the brain CSF and ISF.

If increased serine synthesis in PHGDH-expressing aggressive brain metastatic cells was responsible for the differential proliferation rates observed in CSF and -Ser/-Gly media, suppression of PHGDH in aggressive brain metastatic cells should also inhibit cell growth in a serine- and glycine-limited environment. To demonstrate the necessity of serine synthesis for the phenotypes we observed, we suppressed PHGDH with two independent short hairpin RNAs (shPHGDH #1 and #2; Supplementary Fig. S2E), transduced cells with a control hairpin (shCtrl; either shNonTargeting, shNT or shGFP), and measured these hairpins’ effects on both glucose-derived serine synthesis and cell proliferation. As expected, serine labeling fell substantially by ∼8 – fold in PHGDH-depleted cells, but serine pools remained comparable to wild-type cells in PHGDH-depleted cells grown in plasma concentrations of serine and glycine (Fig. 2E; Supplementary Fig. S2F, G). However, PHGDH-depleted cells had a 30-50% decrease in serine pools when they were grown at CSF serine and glycine concentrations or in -Ser/-Gly media (Supplementary Fig. S2G). These results are consistent with the observation that PHGDH suppression not only abolished glucose-derived serine synthesis, but also impeded aggressive brain metastatic cell proliferation in serine and glycine limited conditions (Fig. 2F). In contrast, aggressive brain metastatic cells cultured in CSF or -Ser/-Gly media, maintained higher serine pools, in which ∼45% of the serine in these pools was derived from glucose (Fig. 2E), and proliferated proficiently in serine and glycine-limited media (Fig. 2F).

To ensure that this phenotype was due to the catalytic activity of PHGDH, we reconstituted PHGDH-depleted cells with catalytically active or inactive shRNA-resistant PHGDH (Supplementary Fig. S2H). Expression of catalytically active shRNA-resistant PHGDH restored glucose-derived serine production and proliferation of PHGDH-depleted aggressive brain metastatic cells in both CSF or -Ser/-Gly media, while expression of an shRNA-resistant, catalytically inactive PHGDH mutant (D175N, R236K, H283A; Supplementary Fig. S2I) failed to restore serine synthesis and cell growth in serine and glycine limited conditions (Fig. 2G, H; Supplementary Fig. S2J, K). Taken together, these results indicate that PHGDH activity promotes glucose-derived serine synthesis and cell proliferation in serine- and glycine-limited conditions, and is an important characteristic of cancer cells that efficiently colonize the brain.

### PHGDH Enables Brain Metastasis

Our *in vitro* data supports the hypothesis that serine synthesis is essential for cancer cell growth in low serine and glycine environments, such as the brain interstitial space. We therefore assessed the necessity of PHGDH for metastasis to the brain *in vivo*. We transduced aggressive brain metastatic cells expressing GFP and luciferase with control (shCtrl) hairpins and two independent hairpins to suppress PHGDH (shPHGDH #1 and shPHGDH #2; Supplementary Fig. S2e), injected these cells into the left ventricle of athymic Nude mice, and monitored metastatic progression by BLI. Mice injected with cells in which PHGDH was suppressed had substantially reduced brain metastasis and improved survival, whereas mice injected with cells transduced with a control shRNA had greater metastatic burden and shorter lifespan (Fig. 3A, B).

**Figure 3.**
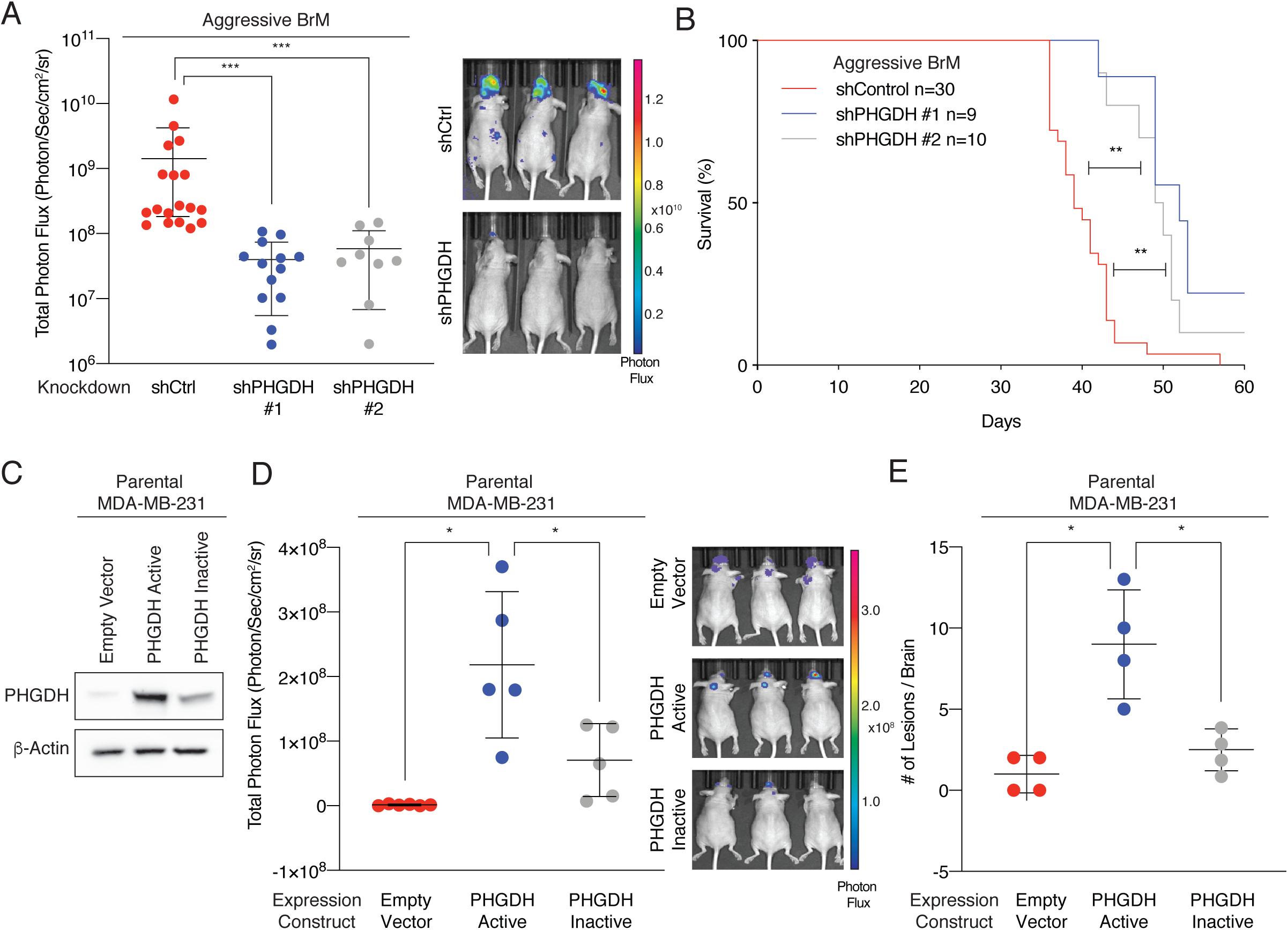
Genetic Depletion of PHGDH Suppresses Brain Metastasis. **(A)** Total photon flux of brain metastasis bearing mice injected with aggressive brain metastatic cells expressing either a control shRNA hairpin or shRNA hairpin targeting PHGDH. PHGDH suppression reduces the brain burden as measured by photon flux of aggressive brain-trophic cells at 28 days post injection. **(B)** Kaplan-Meier plot showing disease-specific survival of mice injected with efficient brain metastatic cells expressing either a control shRNA (shCtrl) or PHGDH shRNA (shPHGDH #1 or shPHGDH #2). Significance was tested using the log-rank test. Suppression of PHGDH in aggressive brain-trophic cells extends the lifespan of mice following intracardiac injection. **(C)** Western blot of parental MDA-MB-231 cells expressing empty-vector (EV), catalytically active or catalytically inactive PHGDH. Parental MDA-MB-231 cells do not express detectable levels of PHGDH but both active and inactive PHGDH are detected in these cells following transduction. **(D)** Brain photon flux of mice injected with parental MDA-MB-231 cells expressing an empty-vector control, catalytically active PHGDH or catalytically inactive PHGDH. Catalytically active PHGDH promotes brain tropism above the empty vector or catalytically inactive PHGDH as measured by photon flux at 28 days post injection. **(E)** Quantitation of brain metastases from parental MDA-MB-231 brain metastases by direct GFP visualization of isolated brains. Catalytically active PHGDH increases the number of lesions per brain in comparison to catalytically inactive PHGDH and transduction with the empty vector.

To determine if PHGDH alone enabled brain metastasis formation, we expressed catalytically active PHGDH or catalytically inactive PHGDH in parental MDA-MB-231 cells, which are not intrinsically brain-trophic (Fig. 3C). Expression of catalytically active PHGDH enabled brain metastasis formation, whereas expression of catalytically inactive PHGDH or transduction of cells with an empty vector did not increase MDA-MB-231 brain metastasis formation greatly following intracardiac injection, as indicated by BLI intensity and quantification of GFP-positive brain metastatic lesions (Fig. 3D, E; Supplementary Fig. S3A). Notably, we did not observe bioluminescence signal in other organs in these experiments, suggesting that PHGDH expression may be dispensable for other tissues excluding the brain. Together, these observations support a model in which PHGDH activity generates serine to enable brain metastasis, and indicate that PHGDH suppression in tumor cells can reduce cancer cell growth in the brain.

### Pharmacological Inhibition of PHGDH Suppresses Cell Growth in Serine- and Glycine-Limited Environments

First generation PHGDH inhibitors can slow the growth of primary PHGDH-dependent tumors, but these tool compounds are neither potent nor selective enough to be useful in humans. Second-generation PHGDH inhibitors have higher potency and improved pharmacokinetic and pharmacodynamic properties, but to date these have not been effective in models of PHGDH-driven malignancies. As our data demonstrate that PHGDH catalytic activity promotes brain metastasis, and the selectivity of cellular responses to genetic suppression of PHGDH, we were encouraged to test second-generation PHGDH inhibitors in our model of brain metastasis.

PH-719 and PH-755 (Raze Therapeutics) are second-generation PHGDH inhibitors with nanomolar potency *in vitro*, are orally bioavailable, and have a >24 hour half-life in mice.^52^ PH-719 completely inhibited serine production, cell proliferation, and invasion in serine- and glycine-limiting cell culture conditions, but did not affect their growth when serine and glycine were at plasma concentrations (Fig. 4A, B; Supplementary Fig. S3B-E). Likewise, PH-755 also inhibited PHGDH in human cells in a concentration-dependent fashion and exhibited a 100-fold improvement in potency over NCT-503, a previously published PHGDH inhibitor (Supplementary Fig. S3F). Moreover, treatment of two PHGDH independent cell lines (MDA-MB-231 and ZR-75-1) and four PHGDH dependent cell lines (HT1080, HCC70, MDA-MB-468, and BT-20) in dose-response assays with PH-719 demonstrated that PHGDH inhibitors had EC_50_ values for cell growth inhibition of 0.5 – 5μM only when these cells were grown in media with CSF concentrations of serine and glycine (Supplementary Fig. S3G, H). PH-719 had a 6 to 10-fold higher EC_50_ for MDA-MB-231 cells, which do not express PHGDH, and no toxicity toward other PHGDH independent cell lines (Supplementary Fig. S3H). Importantly, aggressive brain metastatic cells treated with PHGDH inhibitor showed reduced capacity to synthesize serine and proliferate in serine and glycine limited conditions on par with genetically depleted PHGDH cells (Fig. 4B; Supplementary Fig. S3B, D, E). These results support the hypothesis that extracellular serine availability modulates cellular sensitivity to PHGDH inhibition.

**Figure 4.**
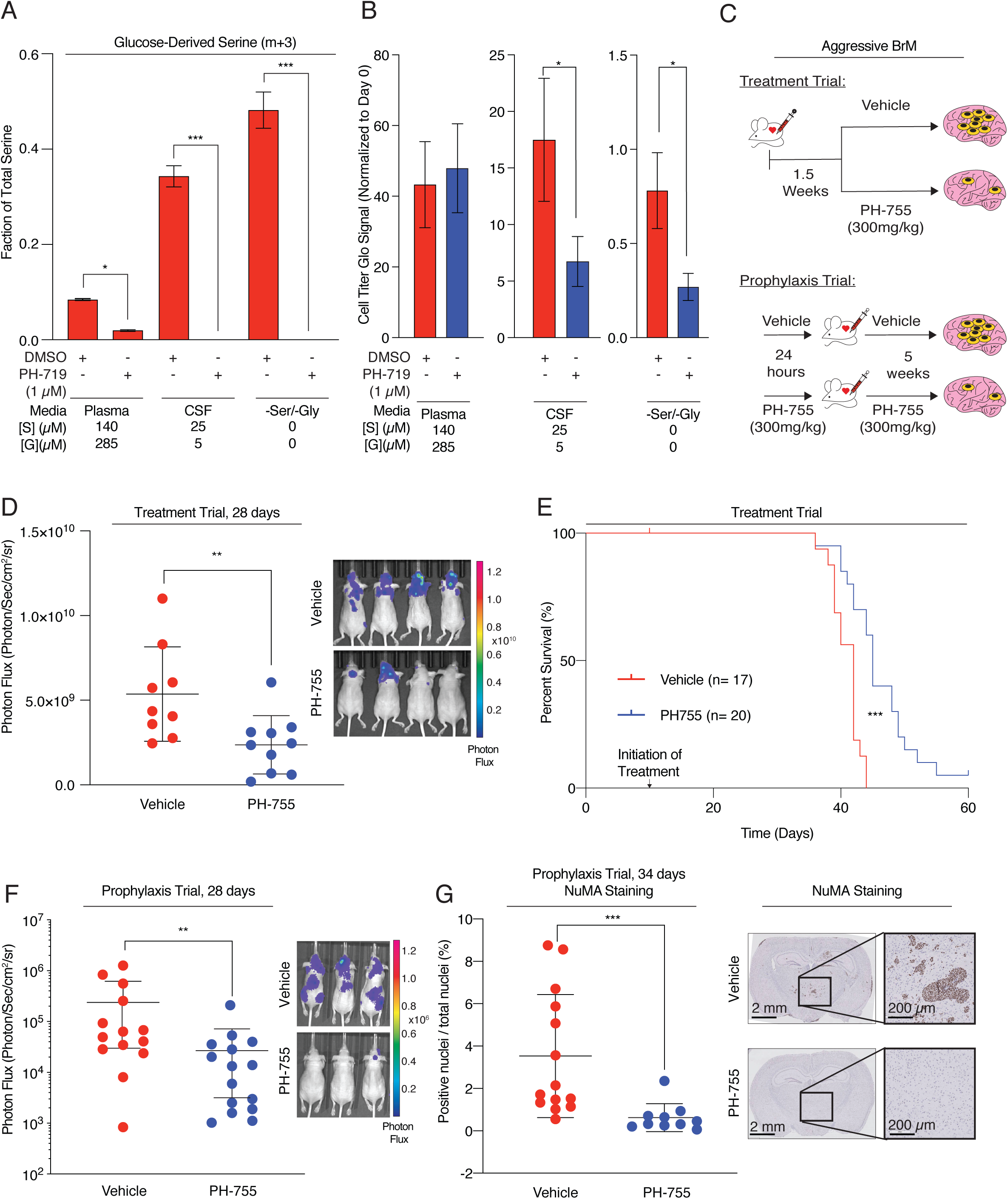
Pharmacological Inhibition of PHGDH Suppresses Brain Metastasis. **(A)** Fractional labeling of intracellular glucose-derived m+3 serine from aggressive brain metastatic cells treated with DMSO or 1 *μ*M of PH-719 in Plasma, CSF, or -Ser/-Gly media. PH-719 blocks the synthesis of glucose-derived serine in aggressive brain-trophic cell lines regardless of extracellular serine and glycine concentrations. **(B)** Proliferative capacity of aggressive brain metastatic cells treated with DMSO or PH719 (1 *μ*M) in Plasma, CSF, or -Ser/Gly media. PH-719 treatment attenuates growth of aggressive brain metastatic cells when cultured in CSF, or -Ser/Gly media. **(B)** Schematic of a mouse clinical trial in which mice were either injected with aggressive brain metastatic cells, and treated 1.5 weeks later with vehicle or 300mg/kg of PH-755 twice daily (treatment trial), or where PH-755 treatment preceded the injection of aggressive brain metastatic cells (prophylaxis trial). **(D)** Total photon flux of brain metastasis-bearing mice treated with vehicle or 300 mg/kg of PH-755 twice daily at week 4. Treatment began 1.5 weeks after the initial introduction of cancer cells into mice. PH-755 treatment reduced brain metastatic burden compared to vehicle control as measured by bioluminescence imaging. **(E)** Kaplan-Meier plot showing disease-specific survival of brain metastasis bearing mice treated with PH-755. Treatment with PH-755 significantly improved the survival of mice bearing brain metastases. Significance tested using log-rank test. **(F)** Total photon flux of brain metastasis bearing mice treated prophylactically with vehicle or 300 mg/kg PH-755 twice daily at week 4. Prophylactic treatment with PH-755 significantly reduces the burden of brain metastases. **(G)** Quantitation of IHC for human Nuclear Mitotic Apparatus (NuMA)-positive nuclei relative to total nuclei count, indicating the burden of brain metastases, from mice treated prophylactically with vehicle or PH-755. The timepoint was 34 days. PH-755 treatment significantly reduces the burden of brain metastases.

### Pharmacological Inhibition of PHGDH Attenuates Brain Metastasis

To evaluate the effect of PH-755 on established brain metastases *in vivo*, we allowed aggressive brain metastatic cells to seed the brains of nude mice, and treated the animals with vehicle or PHGDH inhibitor (300mg/kg twice daily by oral gavage) 1.5 weeks after intracardiac injection when brain metastases were established and visible by BLI (Fig. 4c). Twice-daily dosing of PH-755 ensured continuously high plasma concentrations of the compound.^36^ Treatment with PH-755 for three weeks reduced brain metastatic burden and extended the survival of mice bearing brain metastases (Fig. 4D, E; Supplementary Fig. S4B, C, D). Interestingly, while PH-755 attenuated brain metastases in xenograft models (Fig. 4D, E), it failed to restrict tumor growth of aggressive brain metastatic cells that were implanted into the mammary fat pad (Supplementary Fig. S4A). This highlights the possibility that differences in nutrient availability affect cellular responses to PHGDH inhibition.

Preventing the spread of cancer to the brain would also offer significant benefit to patients with tumors that have a high propensity to spread to the brain.^53^ To test whether prophylactic treatment with PHGDH inhibitors would attenuate brain metastasis formation, we administered PH-755 at 300mg/kg twice daily for 24 hours to athymic nude mice prior to left ventricular inoculation of aggressive brain metastatic cells (Fig. 4C). Over the five-week treatment course, PH-755 significantly reduced metastatic tumor growth and increased metastasis-free survival as measured by bioluminescence imaging compared to vehicle treatment (Fig. 4F; Supplementary Fig. S4E). Histologic examination of the brains of vehicle and PH-755 treated mice revealed a significantly higher burden of brain metastases in the vehicle-treated mice compared to the PH-755 treated mice as determined by automated scoring of human Nuclear Mitotic Apparatus (NuMA) and manual scoring (by two independent observers) of H&E-stained sections (Fig. 4G; Supplementary Fig. S4F). Overall, the number and size of brain metastatic lesions were ∼20 times higher in mice treated with vehicle than in mice treated with PH-755 and no metastases were identified in ∼50% of the PH-755 treatment group (Fig. 4G; Supplementary Fig. S4F).

To determine the pharmacodynamic consequences of treating mice with PH-755, we quantified the concentrations of amino acids in the plasma and CSF of tumor-bearing mice treated with vehicle or drug. PH-755 lowered plasma and CSF serine concentrations, but did not affect the concentrations of other metabolites (Supplementary Fig. S5A, B). U-^13^C-glucose labeling of mice fasted for 6 hours revealed that PH-755 decreased glucose incorporation into serine in both the plasma and brain metastases (Supplementary Fig. S5C-E). Importantly, mice treated with PH-755 did not lose weight during the 8-week treatment in spite of the potential systemic toxicities of inhibiting serine biosynthesis (Supplementary Fig. S5F). Taken together, we believe that this work serves as proof of concept that PHGDH inhibitors might be useful in the treatment and prevention of brain metastases, and support our findings that serine and glycine availability dictates drug response to PHGDH inhibition *in vivo*.

## Discussion

Brain metastasis is a major contributor to cancer mortality, with few therapeutic options. Brain metastases arise in cancers originating from multiple tissues of origin with heterogeneous genetic backgrounds. Therefore, metabolic adaption to the serine- and glycine-depleted metabolic microenvironment of the brain may be a common pathway that tumors require to colonize the brain. We hypothesized that targeting these metabolic dependencies might reduce the initiation or progression of brain metastases. Here, we demonstrate that *de novo* glucose-derived serine synthesis enables brain metastasis formation, and that targeting serine synthesis with small molecules attenuates brain metastasis in mice. These data are consistent with recent evidence suggesting that cancer cells from disparate origins may converge to adopt similar metabolic phenotypes in a given organ.^54-56^

In support of this hypothesis, we demonstrated that expression of catalytically active PHGDH in a non-brain trophic tumor cell line enables brain metastasis, while catalytically inactive PHGDH is far less efficient at promoting brain metastasis. Suppression of PHGDH decreases brain metastatic burden and improves the survival of mice with brain metastasis. Treatment with a second-generation PHGDH inhibitor decreased brain metastatic burden and prolonged survival in mice with established brain metastases. Therefore, we hypothesize that this is due to the serine- and glycine-limited environment of the brain, in which serine concentrations are below the reported K_m_ values for the primary serine transporters ASCT1 and ASCT2, enforces serine synthesis dependencies on cells that enter the brain microenvironment (Supplementary Fig. S5g). Our data also suggest that PHGDH inhibitors may be useful in slowing the progression of established brain metastases in a treatment setting, and potentially in a prophylactic setting, where there is high risk of brain metastasis, particularly for tumors of the breast, lung, kidney, and skin that have a high propensity to spread to the brain.^57^

Although our data demonstrate that brain metastasis formation is enhanced by the catalytic activity of PHGDH, PHGDH may also have non-catalytic functions. As evidence for this hypothesis, we note that aggressive, MDA-MB-231-derived brain metastatic cells appear to tolerate genetic and pharmacological ablation of PHGDH under serine and glycine replete conditions. In contrast, MDA-MB-468 cells are sensitive to genetic suppression of PHGDH in abundant serine conditions, but are unaffected by second generation PHGDH inhibitors unless serine is limiting. Both NCT-503 and CBR-5884, first generation PHGDH inhibitors, have reactive thiol or thiourea groups, which may destabilize the tetrameric, active form of PHGDH, in a manner similar to other thiol reactive drugs, such as disulfiram.^58^ Although second generation PHGDH inhibitors are more potent, they may not mimic PHGDH suppression by destabilizing the tetrameric, active form of the enzyme. This may explain why second generation PHGDH inhibitors are only effective when serine and glycine are limiting, and further emphasizes the necessity of performing rescue experiments with catalytically active and inactive forms of metabolic enzymes in validating a genetic hit as a drug target.

PHGDH inhibitors may be help treat brain metastases, but will inhibit *de novo* serine synthesis in non-malignant cells. Previous studies have shown that PHGDH is highly expressed in radial glial cells, antitumor CD8+ T-cells, astrocytes, macrophages, and endothelial cells.^52,59^ Humans with hypomorphic mutations in PHGDH exhibit developmental delay and mental retardation, both of which can be ameliorated by serine supplementation, suggesting that transitory PHGDH inhibition following completion of neuronal development may be tolerable. While our study found no evidence of neurotoxicity and weight loss due to PHGDH inhibition, future studies will be needed to determine whether PHGDH inhibition needs to be staggered with therapies that require functional PHGDH-dependent cells, such as cytotoxic CD8+ T-lymphocytes.

The development of novel antimetabolites will require a deep understanding of the metabolic vulnerabilities of cancer and the microenvironments that enforce these metabolic dependencies. Our work uses tailored cell culture media and refined mouse models to reveal that brain metastases depend on serine synthesis. The selectivity, tolerability and efficacy of second generation PHGDH inhibitors suggest that these compounds may be useful to attenuate brain metastasis formation and progression.

## Methods

### Knockdown and overexpression constructs

Luciferase expression was achieved using pLVX plasmid (expressing eGFP and luciferase) and cells stably expressing luciferase were sorted for eGFP expression. PHGDH expression was achieved using the pMXS retroviral expression system. Cells were selected using G418 (0.5mg/ml) or puromycin (2ug/ml), respectively. Stable knockdown of human PHGDH was achieved using shRNAs in pLKO.1 as previously described^37^. 2-4 distinct shRNA hairpins were screened per target. Targeted shRNA sequences are listed in Supplementary Table S1. Cell lines were authenticated by STR profiling (Duke DNA Analysis Facility) and tested negative monthly for mycoplasma by PCR.

### Cell Culture and Media

Cells were grown in Dulbecco’s MEM (DMEM) Media without glucose, glutamine, serine, glycine, and sodium pyruvate (US Biological D9802-01) or DMEM media with L-Glutamine and Sodium Bicarbonate, without glucose, serine, and glycine. A 100 x amino acid stock and a non-essential amino acid mix lacking serine and glycine was prepared as a 50x stock and added to the media, along with serine and glycine as needed. Dialyzed inactivated fetal bovine serum (dIFS) was added to the media to 10% (v/v) and the complete media was filtered through a 0.2 μm filter prior to use. Cells were grown in 10 cm dishes in a 5% CO2, ambient oxygen incubator and trypsinzed when confluent (roughly every three days). For growth curves, cells were grown for 4-6 days and counted using an Incucyte (Essen Biosciences) or a Z2 Coulter Counter (Beckman Coulter). Small molecule dose-response curves were printed using an D300e Digital Dispenser (HP Specialty Printing Systems) with DMSO concentrations held at <0.5% of the culture volume and cell growth was measured by Cell Titer Glo (Promega) after 4 days of growth. All plots, curve fitting, and statistical analysis were carried out with GraphPad Prism.

### Proteomic and Phospho-proteomic Profiling

2×10^6^ cells were collected and washed three times with cold PBS before flash-freezing the samples in liquid nitrogen and shipping on dry ice. Control and brain metastases samples were thawed on ice and prepared following the in-stage tip sample preparation method with minor modifications^60^ Briefly, 100 μL of the reducing alkylating sodium deoxycholate buffer (PreOmics, Martinsried, Germany) was added to the samples before protein denaturation at 100 °C for 20 min. Samples were further homogenized by 15-min sonication in a Biorupter (30 s on/off cycles, high settings). Proteins were then digested by Lys-C and trypsin overnight at 37 °C on a shaker set at 1,200RPM. Peptides were acidified to a final concentration of 0.1% trifluoroacetic acid (TFA) for SDB–RPS binding and desalted before LC-MS/MS analysis. Samples were measured on a quadrupole Orbitrap mass spectrometer (Q Exactive HF, Thermo Fisher Scientific, Rockford, IL, USA) coupled to an EASYnLC 1200 ultra-high-pressure system (Thermo Fisher Scientific) via a nanoelectrospray ion source.^61,62^ About 1 μg of peptides was loaded on a 40-cm HPLC-column (75 μm inner diameter; in-house packed using ReproSil-Pur C18-AQ 1.9-μm silica beads; Dr Maisch GmbH, Germany). Peptides were separated using a linear gradient from 3% to 23% B in 82 min and stepped up to 40% in 8 min at 350 nL per min where solvent A was 0.1% formic acid in water and solvent B was 80% acetonitrile and 0.1% formic acid in water. The total duration of the gradient was 100 min. Column temperature was kept at 60 °C by a Peltier element-containing, in-house developed oven. The mass spectrometer was operated in ‘top-15’ data-dependent mode, collecting MS spectra in the Orbitrap mass analyzer (60 000 resolution, 300– 1650 m/z range) with an automatic gain control (AGC) target of 3E6 and a maximum ion injection time of 25 ms. The most intense ions from the full scan were isolated with a width of 1.4 m/z. Following higher-energy collisional dissociation (HCD) with a normalized collision energy (NCE) of 27%, MS/MS spectra were collected in the Orbitrap (15 000 resolution) with an AGC target of 1E5 and a maximum ion injection time of 25 ms. Precursor dynamic exclusion was enabled with a duration of 20 s. Tandem mass spectra were searched against the 2015 Uniprot human databases (UP000005640_9606 and UP000005640_9606_additional) using MaxQuant version 1.5.3.34 with a 1% FDR at the peptide and protein level, peptides with a minimum length of seven amino acids with carbamidomethylation as a fixed modification and protein N-terminal acetylation and methionine oxidations as variable modifications (Cox, J., Mann, M. 2008).^63^ Enzyme specificity was set as C-terminal to arginine and lysine using trypsin as protease, and a maximum of two missed cleavages were allowed in the database search. The maximum initial mass tolerance for precursor and fragment ions was 4.5 ppm and 20 ppm, respectively. If applicable, peptide identifications by MS/MS were transferred between runs to minimize missing values for quantification with a 0.7 min window after retention time alignment. Label-free quantification was performed with the MaxLFQ algorithm using a minimum ratio count of 1. Statistical and bioinformatics analysis was performed with the Perseus software^49^(version 1.5.5.0), Microsoft Excel, and R statistical software. Proteins that were identified in the decoy reverse database or only by site modification were not considered for data analysis. Median of technical triplicates (referring to independent sample preparations) were calculated and mean log_2_ ratios of biological duplicates (two metastases and two control tissues collected at different locations of the pleural cavity), and the corresponding p-values were visualized with volcano plots. We used *t*-tests for binary comparisons and SAM with s0 = 0.1 and a FDR < 0.05 or <0.01 for the assessment of *t*-test results in volcano plots. The FDR was corrected for multiple hypotheses based on permutation-based FDR correction.

### Western Blotting

Tissue and cell lysates were prepared using RIPA buffer (Cell Signaling Technologies) supplemented with protease inhibitor cocktail (Thermo Fisher Scientific). Protein concentrations were quantitated using BioRad Protein Assay. Equal amounts of protein (10-30 μg) were separated on 4-12% Bolt bis-tris plus gels (Thermo Fisher Scientific) and transferred to Immobilon FL PVDF membranes (Millipore Sigma), which were blocked in 5% milk in TBS-T at room temperature for 1 hour and incubated with primary antibodies at 4 °C overnight. Blots were incubated with anti-mouse and anti-rabbit secondary antibodies conjugated to HRP (Cell Signaling Technologies) at room temperature for 1 hour before visualization with Pierce ECL Western Blotting Substrate in a BioRad ChemiDoc Plus Imaging system. Blots were stripped with RestorePlus Western Blot Stripping Buffer (Thermo) prior to probing for the loading control. The following primary antibodies were used: PHGDH (Rabbit Polyclonal; Sigma HPA021241), ERK2 (Rabbit Monoclonal; AbCam ab32081); β-actin (Mouse Monoclonal; Abcam ab8226); HSP90 (Rabbit Polyclonal, Cell Signaling Technologies 4874).

### Metabolite Profiling: Steady-State and Labeling Experiments

Cells were evenly seeded at 250,000 cells per well in a 6-well plate and allowed to attach for 24 hrs. Prior to all labeling experiments, cells were pretreated with 1 μM compound or equivalent volume of DMSO in DMEM for 1 hr. For steady-state metabolite concentrations, cells were washed with PBS before pretreatment and treatment in DMEM lacking serine and glycine. For labeling experiments, U-^13^C-glucose (Cambridge Isotope Laboratories CLM-1396-PK) replaced the corresponding unlabeled glucose DMEM component. Cells were washed in 4 °C 0.9%(w/v) NaCl in LCMS-grade water and extracted in 1mL/well of 3:1:2 methanol:water:chloroform with 500 nM ^13^C, ^15^N-amino acids (Cambridge Isotope Laboratories MSK-A2-1.2) as internal extraction standards. The extraction solvent was dried in a speedvac and metabolite samples were stored at -80C until analysis. Triplicate identically seeded and treated cells were trypsinized and analyzed with a Multisizer Coulter Counter (Beckman Coulter) to obtain cell counts and total cell volumes for normalization.

LC-MS analysis was carried out as described.^36^ In brief, metabolites were resuspended in 100 μL LC-MS grade water and run on a SeQuant ZIC-pHILIC column (2.1×150 mm, 5 μm, Millipore-Sigma) on a Dionex UltiMate 3000 UPLC using a 0 to 80% gradient of 20 mM ammonium carbonate and 0.1% ammonium hydroxide, pH 9, in water. Mobile phase was introduced into the ionization source of a high-field Q Exactive Mass Spectrometer (QE HF; Thermo Fisher Scientific) running in polarity switching mode and identified by retention time, accurate mass, and fragmentation. Metabolite peaks were identified and integrated with Skeleton (NYU Metabolomics Core Resource Laboratory) and normalized to internal standards and total cell volume. m/z ratios for ^13^C -labeled metabolites were calculated using Skeleton and IsoMETLIN and corrected for isotopic abundance.^64^

GC-MS analysis was carried out and analyzed as described.^65^ In brief, dried, extracted metabolites were derivatized using a MOX-tBDMCS method and analyzed by GC-MS using a DB-35MS column (30 m × 0.25 mm i.d., 0.25 μm) in an Agilent 7890B gas chromatograph interfaced with a 5977B mass spectrometer. Metabolites were identified by unique fragments and retention time in comparison to known standards. Peaks were picked in OpenChrom and integrated and corrected for natural isotopic abundance using in-house algorithms adapted from Fernandez et al.^66,67^

### Animal studies

Animal experiments were performed in accordance with protocols approved by the Weill Cornell Medicine and New York University Langone Health Institutional Animal Care and Use Committee (IACUC; Weill Cornell protocol 2013-0116; NYULMC protocol IA16-01862). For knockdown and inhibitor experiments, the minimum group size per arm was 12 for an anticipated difference of 55% between arms assuming a type I error probability of 0.05 and a power of 80%. We used 15 animals per arm in case of animal loss during intracardiac injection or subsequent procedures.

Intracardiac injection was performed as previously described.^1^ Briefly, cells were trypsinized and washed with PBS and a 1×10^5^ cells (in 100 μl of PBS) were injected into the left cardiac ventricle of female 5-6 week old athymic nude (nu/nu) mice (Jackson Laboratory strain 002019). Mice were then immediately injected with D-luciferin (150 mg/kg^-1^) and subjected to bioluminescence imaging (BLI) using the IVIS Spectrum Xenogen instrument (Caliper Life Sciences) to ensure systemic dissemination of tumor cells. Metastatic burden was measured weekly after injection using BLI for up to 17 weeks. BLI images were analyzed using Living Image Software v.2.50. Disease-specific survival endpoints were met when the mice died or met the criteria for euthanasia under the IACUC protocol and had radiographic evidence of metastatic disease. T2 MR Imaging (0.5 mm slices) was obtained using a Bruker Biospec 94/30 USR 9.4 Tesla Small Animal MRI with a B-GA 20S gradient system in the MSKCC Rodent Imaging Core.

For orthotopic tumor implantation, 2.5×10^5^ cells in 100ul of PBS were mixed 1:1 with Matrigel (BD Biosciences) and injected into the fourth mammary fat pad. Only one tumor was implanted per animal. Primary tumors were surgically excited when they reached ∼1.5 cm in the largest dimension and metastatic dissemination was assessed using BLI imaging at 1-3 week intervals for up to 30 weeks. Distant metastasis-free survival endpoint was met when BLI signal was seen outside of site of primary tumor transplantation.

To derive short-term cultures from primary tumors and metastases, anesthetized animals (isoflurane) were imaged then sacrificed. *Ex-vivo* BLI was subsequently performed on harvested organs to define the precise location of the metastatic lesion. Primary tumors and metastases were subsequently mechanically dissociated and cultured in DMEM and sorted for eGFP to exclude host cells.

PH-755 was administered to mice to a dose of 300 mg/kg twice daily by oral gavage as a suspension in PBS containing 0.5% methylcellulose and 0.5% Tween 80. For brain histology, mice were deeply anesthetized with isoflurane and perfused with 10% formalin in PBS prior to sacrifice. Brains were harvested and fixed for 24 hours in 10% formalin followed by 48 hours in 70% ethanol, sectioned, and embedded for H&E staining. Data represent the mean of two independent observers using FIJI to measure brain and metastasis size.

CSF was isolated as previously described.^14^ Whole blood was obtained by cardiac puncture into heparinized collection tubes (BD Microtainer), which were centrifuged at 1000 × g at 4 °C for 10 minutes. The plasma was flash-frozen until analysis. For LC-MS analysis, 20 μL of plasma were extracted with 80 μL 80:20 methanol:water with 500 nM ^13^C, ^15^N-amino acids (Cambridge Isotope Laboratories MSK-A2-1.2) as internal extraction standards and centrifuged at ≥15000 × g at 4 °C for 10 minutes. The supernatant was dried in a speedvac and stored at -80 °C until GC-MS or LC-MS analysis.

Differences in luminescence and in metastasis area were analyzed using the Mann-Whitey *U* test. Kaplan-Meier survival curves were compared using the log-rank test.

For *in vivo* labeling, mice were fasted for 6 hours starting from midnight and injected intraperitoneally with 2 g/kg of U-^13^C-glucose and sacrificed 60 minutes later. Following cardiac puncture to recover plasma, the brain was removed and visualized using a Leica M205 FA fluorescence stereo dissecting scope and metastases dissected quickly using a scalpel and forceps. Tissues were quenched at liquid nitrogen temperatures using a BioSqueezer (BioSpec Products) and stored at -80 °C. Tissue samples were weighed, extracted in 200 μL 3:1:2 methanol:water:chloroform / mg of tissue, and the polar fraction was recovered, dried on a speedvac, and stored at -80 °C until LC-MS analysis or derivatization for GC-MS analysis.

Pharmacokinetic data were obtained from groups of n≥3 mice that were administered PH-755 by oral gavage in suspension at the indicated doses and whole blood drawn and plasma isolated at the indicated timepoints. PH-755 was quantitated by LC-MS using a standard curve.

### Immunohistochemistry

Immunohistochemistry was performed on 10% neutral buffered formalin fixed, paraffin-embedded human and murine tissue sections. Sections were collected at 5-microns onto plus slides (Fisher Scientific, Cat # 22-042-924) and stored at room temperature prior to use. Unconjugated, polyclonal rabbit anti-human Phosphoglycerate Dehydrogenase (Sigma Aldrich Cat# HPA021241 Lot# GR268490-13 RRID: AB_10680001) was used for human tissues. Unconjugated, polyclonal rabbit anti-human Nuclear Mitotic Apparatus Protein (Abcam Cat# 97585 Lot# B115626 RRID: AB_1855299) was used for murine xenograft samples.

Chromogenic Immunohistochemistry was performed on a Ventana Medical Systems Discovery XT instrument with online deparaffinization and using Ventana’s reagents and detection kits unless otherwise noted. Sections for Phosphoglycerate Dehydrogenase labeling were antigen retrieved in Ventana Cell Conditioner 2 (Citrate pH6.0) at 95° C for 20 minutes. Sections for Nuclear Mitotic Apparatus Protein were deparaffinized in xylene and rehydrated in graded ethanol followed by rinsing in deionized water. Epitope retrieval was performed in a 1200-Watt microwave oven at 100% power in 10 mM sodium citrate buffer, pH 6.0 for 10 minutes. Slides were allowed to cool for 30 minutes, rinsed in distilled water, and loaded onto the instrument. Endogenous peroxidase activity was blocked with 3% hydrogen peroxide for 4 minutes. Phosphoglycerate Dehydrogenase antibody was diluted 1:100 in Ventana diluent (Ventan Discovery Antibody Diluent, cat# 760-108) and incubated for 3 hours at room temperature. Nuclear Mitotic Apparatus Protein was diluted 1:7000 in Tris-BSA (25 mM Tris, 15 mM NaCL, 1% BSA, pH 7.2) and incubated for 12 hours. Both antibodies were detected with goat anti-rabbit HRP conjugated multimer for 8 minutes. The immune complexes were visualized with 3,3 diaminobenzidene and enhanced with copper sulfate. Slides were washed in distilled water, counterstained with hematoxylin, dehydrated and mounted with permanent media. Appropriate positive and negative controls were run in parallel to study sections. Imaging was carried out with a Hamamatsu NanoZoomer whole-slide scanner.

For PHGDH scoring, the same board-certified pathologist (M. Snuderl) who scored the specimens in ^32^ scored PHGDH specimens in a blinded fashion for the intensity of PHGDH staining using a scale of 0-3, representing none, weak, moderate, and strong staining. The use of the tumor samples was approved by the NYULMC IRB (protocol S16-00122). Data were analyzed using Fisher’s exact test. NuMA staining was quantitated by automated counting of positive and total nuclei using Visiopharm Image Analysis Software.

### *In Vitro* Invasion Assay

For the invasion and migration assays we used the cytoselect cell invasion (CBA-110) kits. In brief, 3×10^5^ cells were suspended in serum-free medium and placed on top of the membrane. Medium containing serum was placed at the bottom and cells that had invaded to the inferior surface of the collagen membrane were stained and counted 24 hrs. later.

### Isolation of Patient-Derived CTCs

Peripheral blood from metastatic breast cancer patients were processed by CTC-iChip. In brief, CD45/CD15 conjugated magnetic beads were used to deplete white blood cells, and antibody agnostic circulating tumor cells were collected in the effluent. These CTCs were cultured in 4% oxygen, in RPMI with bFGF, EGF, and B27 supplementation^69^. Once CTCs proliferated to have enough biological material for in vivo work, they were transduced with a GFP-luciferase expressing construct and injected by intracardiac injection into NSG mice^70^. These mice were followed for 5-8 months for metastatic relapse by bioluminescent imaging. At endpoint, metastatic lesions were resected and sorted for GFP+ cells. To obtain second generation and third generation brain metastatic derivatives, cells were expanded in culture, and additional rounds of in vivo injection (∼100K cells/mouse) was performed.

### In vitro enzyme activity

The D175N, R236K, H283A catalytically inactive mutant of PHGDH was constructed by overlapping PCR. N-terminally 6xHis tagged wild-type and catalytically inactive PHGDH were expressed, purified, and assayed as described.^36^

## Additional Information

### Financial Support

**B.N.** is supported by the National Science Foundation (NSF) Graduate Research Fellowship and the National Cancer Institute (NCI) of the National Institutes of Health (NIH) under the F99/K00 Career Transition Fellowship (F99CA234950). **A.L.** and **S.M.D.** were supported by an NSF Graduate Research Fellowship and T32GM007287. **A.A.** is supported by the HHMI medical fellows program. **R.E.** is supported by a Tow Foundation Postdoctoral Fellowship from Center for Molecular Imaging and Nanotechnology (CMINT) at Memorial Sloan Kettering. **M.M.** is supported by the Max Planck Society for the Advancement of Science, the European Union’s Horizon 2020 research and innovation program (Grant agreement no. 686547; MSmed project), the European Commission 7^th^ Research Framework Program (GA ERC-202120Syg_318987; ToPAG project), and by the Novo Nordisk Foundation for the Clinical Proteomics group (grant NNF15CC0001). **S.F.B.** is supported by the NIH (DP5OD026395 High-Risk High-Reward Program), the Department of Defense Breast Cancer Research Breakthrough Award W81XWH-16-1-0315 (Project: BC151244), the Burroughs Wellcome Fund Career Award for Medical Scientists, the Parker Institute for Immunotherapy at MSKCC, the Josie Robertson Foundation, and the MSKCC core grant P30-CA008748. **D.M.S.** is supported by the NCI of the NIH under Award RO3 DA034602-01A1, R01 CA129105, R01 CA103866, and R37 AI047389. **D.M.S.** is an investigator of the Howard Hughes Medical Institute. **R.K.J.** and **M.G.V.H.** were supported by a Dana-Farber Harvard Cancer Center/MIT Koch Institute Bridge Project grant. **R.K.J.** is also supported by the Lustgarten Foundation; a grant from the Ludwig Center at Harvard; National Cancer Institute Grants P01-CA080124, R01-CA126642, R01-CA085140, R01-CA115767, R01-CA098706, R01CA208205, U01CA224173, and R35CA197743; and US Department of Defense Breast Cancer Research Program Innovator Award W81XWH-10-1-0016. **M.G.V.H.** is also supported by the NIH (R21CA198028, R01CA168653), the MIT Ludwig Center, the MIT Center for Precision Cancer Medicine, SU2C, the Lustgarten Foundation, and a Faculty Scholar Grant from HHMI. **E.H.** is supported by National Institutes of Health (NIH)/ National Cancer Institute (NCI) R01CA2022027. **M.Y.** is supported by an NCI K22 Career Transition Award (K22 CA175228-01A1), NIH DP2 award (DP2 CA206653), Donald E. And Delia B. Baxter Foundation, Stop Cancer Foundation, Wright Foundation, and the Pew Charitable Trusts, and a Pew-Stewart Scholar for Cancer Research. **L.C.C.** is supported by the NIH (NCI R35CA197588), the Breast Cancer Research Foundation, the Gray Foundation Basser Initiative, and a gift from the Lori and Zachary Schreiber Family. **M.E.P.** is supported by a Mary Kay Foundation Cancer Research Grant (017-32), the Shifrin-Myers Breast Cancer Discovery Fund at NYULMC, a V Foundation V Scholar Grant funded by the Hearst Foundation (V2017-004), an NCI K22 Career Transition Award (1K22CA212059), and laboratory-directed research funding from the John and Elaine Kanas Family Foundation.

## Acknowledgements

The authors would like to thank all members of the Mann, Cantley, and Pacold Laboratories for critical feedback. We also thank Miriam Sindelar, Steven Gross, Lin Lin, and Costas Lyssiotis for technical advice and the NYULH, WCMC and MSK Small Animal Imaging Cores for assistance with MRI and BLI. We are grateful for support from Luis Chiriboga and the NYU Center for Bio-specimen Research and Development, which is partially funded by the NYUPCC Cancer Support Grant, NIH/NCI 5P30CA016087-33, and the NYU Metabolomics Core Resource Laboratory.

## Author Contributions

B.N., L.C.C., and M.E.P. conceived of the project, B.N., E.K., S.D., S.B., A.P., D.K., S.B., R.K.S.,, S.F.B., E.M., D.J., and M.E.P. performed the experiments, B.N., E.K., S.D., S.B., A.P., D.K., S.B., L.C.C., M.M., M.Y., M.S. and M.E.P. assisted with the design and data interpretation. All authors contributed to the writing and editing of the manuscript.

## References

1. Bakhoum, S. F. et al. Chromosomal instability drives metastasis through a cytosolic DNA response. Nature 376, 2109 (2018).

2. McDonald, O. G. et al. Epigenomic reprogramming during pancreatic cancer progression links anabolic glucose metabolism to distant metastasis. Nat Genet 49, 367–376 (2017).

3. Jacob, L. S. et al. Metastatic Competence Can Emerge with Selection of Preexisting Oncogenic Alleles without a Need of New Mutations. Cancer Research 75, 3713–3719 (2015).

4. Valiente, M. et al. The Evolving Landscape of Brain Metastasis. Trends Cancer 4, 176–196 (2018).

5. Kodack, D. P., Askoxylakis, V., Ferraro, G. B., Fukumura, D. & Jain, R. K. Emerging strategies for treating brain metastases from breast cancer. Cancer Cell 27, 163–175 (2015).

6. Achrol, A. S. et al. Brain metastases. Nat Rev Dis Primers 5, 5 (2019).

7. Ostrom, Q. T., Wright, C. H. & Barnholtz-Sloan, J. S. Brain metastases: epidemiology. Handb Clin Neurol 149, 27–42 (2018).

8. Tawbi, H. A. et al. Combined Nivolumab and Ipilimumab in Melanoma Metastatic to the Brain. N. Engl. J. Med. 379, 722–730 (2018).

9. Chen, Q. et al. Carcinoma-astrocyte gap junctions promote brain metastasis by cGAMP transfer. Nature 533, 493–498 (2016).

10. Bos, P. D. et al. Genes that mediate breast cancer metastasis to the brain. Nature 459, 1005–1009 (2009).

11. Priego, N. et al. STAT3 labels a subpopulation of reactive astrocytes required for brain metastasis. Nat. Med. 24, 1024–1035 (2018).

12. Chen, J. et al. Gain of glucose-independent growth upon metastasis of breast cancer cells to the brain. Cancer Research 75, 554–565 (2015).

13. Brastianos, P. K. et al. Genomic Characterization of Brain Metastases Reveals Branched Evolution and Potential Therapeutic Targets. Cancer Discov 5, 1164–1177 (2015).

14. Boire, A. et al. Complement Component 3 Adapts the Cerebrospinal Fluid for Leptomeningeal Metastasis. Cell 168, 1101–1113.e13 (2017).

15. Fischer, G. M. et al. Molecular Profiling Reveals Unique Immune and Metabolic Features of Melanoma Brain Metastases. Cancer Discov (2019). doi: 10.1158/2159-8290.CD-18-1489

16. Er, E. E. et al. Pericyte-like spreading by disseminated cancer cells activates YAP and MRTF for metastatic colonization. Nat. Cell Biol. 20, 966–978 (2018).

17. Valiente, M. et al. Serpins promote cancer cell survival and vascular co-option in brain metastasis. Cell 156, 1002–1016 (2014).

18. Sullivan, M. R. & Vander Heiden, M. G. Determinants of nutrient limitation in cancer. Crit. Rev. Biochem. Mol. Biol. 282, 1–15 (2019).

19. Muir, A. & Vander Heiden, M. G. The nutrient environment affects therapy. Science 360, 962–963 (2018).

20. Sullivan, M. R. & Vander Heiden, M. G. When cancer needs what’s non-essential. Nat. Cell Biol. 19, 418–420 (2017).

21. Garcia-Bermudez, J. et al. Aspartate is a limiting metabolite for cancer cell proliferation under hypoxia and in tumours. Nat. Cell Biol. 20, 755–781 (2018).

22. Davidson, S. M. et al. Direct evidence for cancer-cell-autonomous extracellular protein catabolism in pancreatic tumors. Nat. Med. 23, 235–241 (2017).

23. Gui, D. Y. et al. Environment Dictates Dependence on Mitochondrial Complex I for NAD+ and Aspartate Production and Determines Cancer Cell Sensitivity to Metformin. Cell Metab. 24, 716–727 (2016).

24. Alkan, H. F. et al. Cytosolic Aspartate Availability Determines Cell Survival When Glutamine Is Limiting. Cell Metab. (2018). doi: 10.1016/j.cmet.2018.07.021

25. Muir, A. et al. Environmental cystine drives glutamine anaplerosis and sensitizes cancer cells to glutaminase inhibition. Elife 6, (2017).

26. Sullivan, M. R. et al. Increased Serine Synthesis Provides an Advantage for Tumors Arising in Tissues Where Serine Levels Are Limiting. Cell Metab. (2019). doi: 10.1016/j.cmet.2019.02.015

27. Sullivan, M. R. et al. Quantification of microenvironmental metabolites in murine cancers reveals determinants of tumor nutrient availability. Elife 8, (2019).

28. Cantor, J. R. et al. Physiologic Medium Rewires Cellular Metabolism and Reveals Uric Acid as an Endogenous Inhibitor of UMP Synthase. Cell 169, 258–272.e17 (2017).

29. Davidson, S. M. et al. Environment Impacts the Metabolic Dependencies of Ras-Driven Non-Small Cell Lung Cancer. Cell Metab. 23, 517–528 (2016).

30. Maggs, D. G. et al. Interstitial fluid concentrations of glycerol, glucose, and amino acids in human quadricep muscle and adipose tissue. Evidence for significant lipolysis in skeletal muscle. J Clin Invest 96, 370–377 (1995).

31. Commisso, C. et al. Macropinocytosis of protein is an amino acid supply route in Ras-transformed cells. Nature 497, 633–637 (2013).

32. Possemato, R. et al. Functional genomics reveal that the serine synthesis pathway is essential in breast cancer. Nature 476, 346–350 (2011).

33. Locasale, J. W. et al. Phosphoglycerate dehydrogenase diverts glycolytic flux and contributes to oncogenesis. Nat Genet 43, 869–874 (2011).

34. DeNicola, G. M. et al. Oncogene-induced Nrf2 transcription promotes ROS detoxification and tumorigenesis. Nature 475, 106–109 (2011).

35. Zhang, B. et al. PHGDH Defines a Metabolic Subtype in Lung Adenocarcinomas with Poor Prognosis. Cell Rep 19, 2289–2303 (2017).

36. Pacold, M. E. et al. A PHGDH inhibitor reveals coordination of serine synthesis and one-carbon unit fate. Nat. Chem. Biol. 12, 452–458 (2016).

37. Mullarky, E. et al. Identification of a small molecule inhibitor of 3-phosphoglycerate dehydrogenase to target serine biosynthesis in cancers. Proc. Natl. Acad. Sci. U.S.A. 113, 1778–1783 (2016).

38. Reid, M. A. et al. Serine synthesis through PHGDH coordinates nucleotide levels by maintaining central carbon metabolism. Nat Commun 9, 5442 (2018).

39. Chen, J. et al. Phosphoglycerate dehydrogenase is dispensable for breast tumor maintenance and growth. Oncotarget 4, 2502–2511 (2013).

40. Valiente, M. et al. The Evolving Landscape of Brain Metastasis. Trends Cancer 4, 176–196 (2018).

41. Geiger, T., Cox, J. & Mann, M. Proteomic changes resulting from gene copy number variations in cancer cells. PLoS Genet. 6, e1001090 (2010).

42. Zhang, B. et al. Proteogenomic characterization of human colon and rectal cancer. Nature 513, 382–387 (2014).

43. Schwanhäusser, B. et al. Global quantification of mammalian gene expression control. Nature 473, 337–342 (2011).

44. Coscia, F. et al. Multi-level Proteomics Identifies CT45 as a Chemosensitivity Mediator and Immunotherapy Target in Ovarian Cancer. Cell 175, 159–170.e16 (2018).

45. Eckert, M. A. et al. Proteomics reveals NNMT as a master metabolic regulator of cancer-associated fibroblasts. Nature 569, 723–728 (2019).

46. Doll, S., Gnad, F. & Mann, M. The Case for Proteomics and Phospho-Proteomics in Personalized Cancer Medicine. Proteomics Clin Appl 13, e1800113 (2019).

47. Doll, S. et al. Rapid proteomic analysis for solid tumors reveals LSD1 as a drug target in an end-stage cancer patient. Mol Oncol 12, 1296–1307 (2018).

48. Mattaini, K. R., Sullivan, M. R. & Vander Heiden, M. G. The importance of serine metabolism in cancer. J. Cell Biol. 214, 249–257 (2016).

49. Tyanova, S. et al. Proteomic maps of breast cancer subtypes. Nat Commun 7, 10259 (2016).

50. Kit, S. The biosynthesis of free glycine and serine by tumors. Cancer Research 15, 715–718 (1955).

51. Dolgodilina, E. et al. Brain interstitial fluid glutamine homeostasis is controlled by blood-brain barrier SLC7A5/LAT1 amino acid transporter. J. Cereb. Blood Flow Metab. 36, 1929–1941 (2016).

52. Rodriguez, A. E. et al. Serine Metabolism Supports Macrophage IL-1β Production. Cell Metab. 29, 1003–1011.e4 (2019).

53. Slotman, B. et al. Prophylactic cranial irradiation in extensive small-cell lung cancer. N. Engl. J. Med. 357, 664–672 (2007).

54. Schild, T., Low, V., Blenis, J. & Gomes, A. P. Unique Metabolic Adaptations Dictate Distal Organ-Specific Metastatic Colonization. Cancer Cell 33, 347–354 (2018).

55. Basnet, H. et al. Flura-seq identifies organ-specific metabolic adaptations during early metastatic colonization. Elife 8, 5978 (2019).

56. Mashimo, T. et al. Acetate is a bioenergetic substrate for human glioblastoma and brain metastases. Cell 159, 1603–1614 (2014).

57. Aupérin, A. et al. Prophylactic cranial irradiation for patients with small-cell lung cancer in complete remission. Prophylactic Cranial Irradiation Overview Collaborative Group. New England Journal of Medicine (USA) 341, 476–484 (1999).

58. Spillier, Q. et al. Anti-alcohol abuse drug disulfiram inhibits human PHGDH via disruption of its active tetrameric form through a specific cysteine oxidation. Sci Rep 9, 4737–9 (2019).

59. Vandekeere, S. et al. Serine Synthesis via PHGDH Is Essential for Heme Production in Endothelial Cells. Cell Metab. (2018). doi: 10.1016/j.cmet.2018.06.009

60. Kulak, N. A., Pichler, G., Paron, I., Nagaraj, N. & Mann, M. Minimal, encapsulated proteomic-sample processing applied to copy-number estimation in eukaryotic cells. Nat. Methods 11, 319–324 (2014).

61. Kelstrup, C. D. et al. Rapid and deep proteomes by faster sequencing on a benchtop quadrupole ultra-high-field Orbitrap mass spectrometer. J. Proteome Res. 13, 6187–6195 (2014).

62. Scheltema, R. A. et al. The Q Exactive HF, a Benchtop mass spectrometer with a pre-filter, high-performance quadrupole and an ultra-high-field Orbitrap analyzer. Mol. Cell Proteomics 13, 3698–3708 (2014).

63. Cox, J. & Mann, M. MaxQuant enables high peptide identification rates, individualized p.p.b.-range mass accuracies and proteome-wide protein quantification. Nat. Biotechnol. 26, 1367–1372 (2008).

64. Cho, K. et al. isoMETLIN: a database for isotope-based metabolomics. Anal. Chem. 86, 9358–9361 (2014).

65. Parker, S. J. et al. LKB1 promotes metabolic flexibility in response to energy stress. Metab. Eng. 43, 208–217 (2017).

66. Wenig, P. & Odermatt, J. OpenChrom: a cross-platform open source software for the mass spectrometric analysis of chromatographic data. BMC Bioinformatics 11, 405–9 (2010).

67. Fernandez, C. A., Rosiers, Des C., Previs, S. F., David, F. & Brunengraber, H. Correction of 13C mass isotopomer distributions for natural stable isotope abundance. J Mass Spectrom 31, 255–262 (1996).

68. Rainesalo S., Keränen T., Palmio J., Peltola J., Oja S. S., & Saransaari P. Plasma and cerebrospinal fluid amino acids in normal and epileptic patients. Neurochem. Res. 29, 319–324 (2004).

69. Yu, M. et al. Cancer therapy. Ex vivo culture of circulating breast tumor cells for individualized testing of drug susceptibility. Science 345, 216–220 (2014).

70. Klotz, R. et al. Circulating tumor cells exhibit metastatic tropism and reveal brain metastasis drivers. Cancer Discov CD–19–0384 (2019). doi: 10.1158/2159-8290.CD-19-0384

