## Supplementary Figures S1-S5, Supplementary Table S1 for "Limited Environmental Serine Confers Sensitivity to PHGDH Inhibition in Brain Metastasis"

### Supplementary Figure S1:

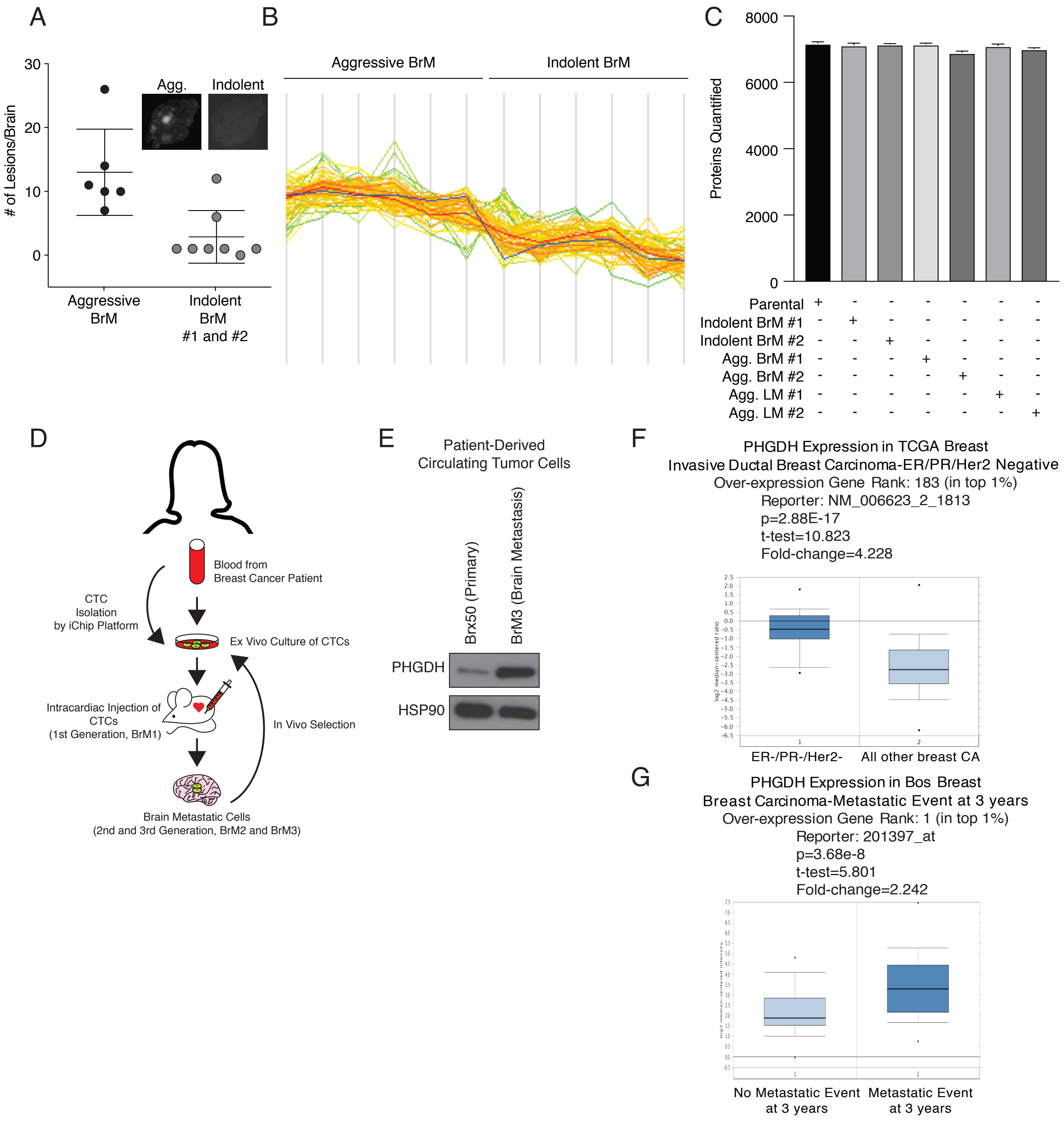

**Supplementary Figure S1, Related to Figure 1.**

**(A)** Number of aggressive and indolent metastatic lesions per brain (Brain metastases; BrM) quantified using a fluorescent dissection microscope. Indolent brain-trophic cells form fewer tumors than aggressive brain-trophic cells. **(B)** PHGDH Peptide intensity in aggressive and indolent brain metastatic derivatives (blue). The number of peptides is higher in aggressive BrM cells compared to indolent BrM cells **(C)** Number of proteins quantified per sample. The number of proteins is similar in all groups. **(D)** Schematic of CTC isolation from breast cancer patient. **(E)** Western blot analysis of PHGDH expression in parental patient derived circulating tumor cells and CTCs that metastasize to the brain. PHGDH is higher in brain-trophic CTCs. **(F)** Oncomine expression data comparing TCGA breast cancer PHGDH expression levels, showing that PHGDH is higher in triple-negative breast cancers compared to non-metastatic tumors. **(G)** Oncomine expression data of PHGDH expression levels in primary breast cancers and metastases. PHGDH is significantly higher at the transcript level in patients with a metastatic event compared to patients who did not have metastases at 3 years.

### Supplementary Figure S2:

A

| Mouse | [Amino Acid] (μM) | |
|---|---|---|
| | Serine | Glycine |
| Plasma | 109 | 246 |
| CSF | 37 | 34 |
| Brain ISF | 9.8 | 7.8 |

See legend for references

B

Working concentrations

| Media | [Amino Acid] (μM) | |
|---|---|---|
| | Serine | Glycine |
| Plasma | 140 | 280 |
| CSF | 25 | 5 |
| -Ser/-Gly | 0 | 0 |

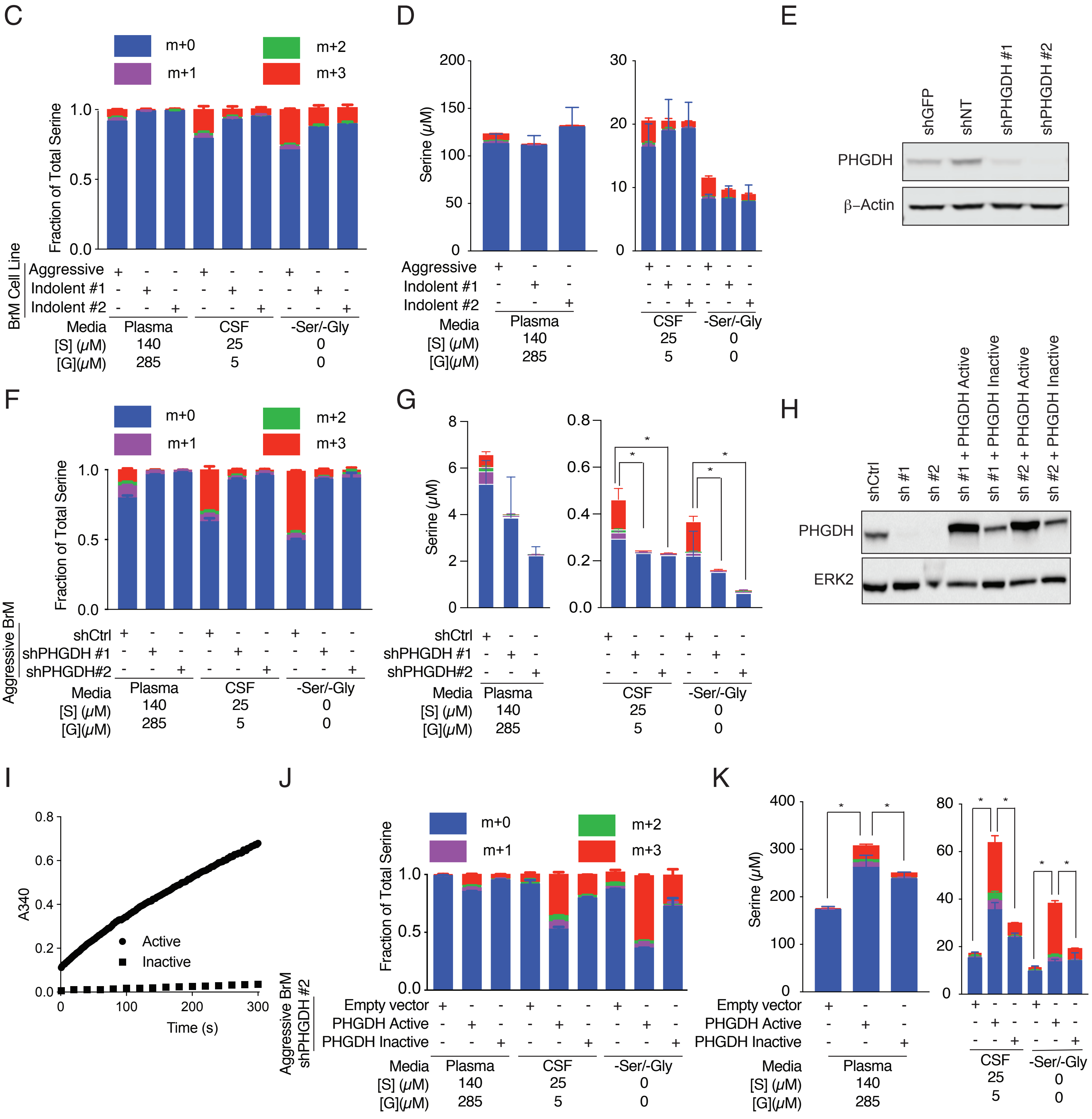

**Supplementary Figure S2, Related to Figure 2.**

**(A)** Published concentrations of serine and glycine in mouse plasma, cerebrospinal fluid (CSF), and the brain interstitial fluid (ISF)[30,51]. The CSF and ISF concentrations are comparable and substantially lower than plasma concentrations.
**(B)** Concentrations of serine and glycine in Plasma, CSF, and -Ser/-Gly cell culture media. These are based on [68] and our measurements (Fig. 2b). **(C)**. Fractional labeling of U-13C-glucose-derived serine from aggressive brain metastatic cells and indolent brain metastatic cells cultured in plasma, CSF, and -Ser/-Gly media. Aggressive BrM cells display enhanced capacity to synthesize serine compared to Indolent BrM cells. **(D)** Intracellular concentrations of unlabeled, exogenous serine and m+3, glucose-derived serine in aggressive and indolent brain metastatic cells in media with plasma, CSF, and null serine and glycine. The concentrations of serine are lower in serine and glycine-low or -null media and the fraction of glucose-derived serine is higher. Indolent cell lines make less glucose-derived serine in comparison to aggressive brain-trophic cells.
**(E)** Western blot of PHGDH expression in aggressive brain metastatic cells transduced with control shRNAs (shGFP and shNT; further labeled as shCtrl) or shRNAs targeting PHGDH (shPHGDH #1 and shPHGDH #2). Both hairpins targeting PHGDH suppress PHGDH expression at the protein level. **(F)** Fractional labeling of U-13C-glucose derived serine from aggressive brain metastatic cells transduced with shCtrl or shRNAs targeting PHGDH (shPHGDH #1 or shPHGDH #2) and cultured in plasma, CSF, or -Ser/-Gly media. Genetic suppression of PHGDH attenuates the production of m+3, glucose-derived serine. **(G)** Total serine pools, normalized to cell counts, from fractional labeling of U-13C- glucose derived serine from aggressive brain metastatic cells depleted of PHGDH and cultured in plasma, CSF, and -Ser/-Gly media. As in Supplementary Fig. 2f, PHGDH suppression abrogates the production of m+3, glucose-derived serine. **(H)** Western blot of aggressive brain metastatic cells depleted of PHGDH, expressing empty vector, catalytically active PHGDH, or catalytically inactive PHGDH. PHGDH knockdown suppresses detectable PHGDH at the protein level, while expression of catalytically active or catalytically inactive PHGDH results in detectable expression of the enzyme. **(I)** *In vitro* enzymatic activity assay of catalytically active PHGDH and inactive PHGDH (D175N, R236K, H283A). The inactive PHGDH does not generate NADH as measured by absorbance at 340 nm. **(J)** Fractional labeling of U-13C-glucose derived serine from aggressive brain metastatic cells depleted of PHGDH, and expressing an empty vector (EV) control, RNAi resistant, codon optimized catalytically active (wildtype) or inactive (D175N, R236K, H283A) PHGDH. Cells were cultured in plasma, CSF, and -Ser/-Gly media. Rescue with catalytically active PHGDH restores serine synthesis, whereas catalytically inactive PHGDH is unable to completely rescue serine synthesis. **(K)** Serine pools, normalized to cell volume, in aggressive brain metastatic cells with suppressed PHGDH with addback of catalytically active and inactive PHGDH. Active PHGDH restores production of m+3, glucose-derived serine, while the empty vector or catalytically inactive PHGDH fail to do so.

Supplementary Figure S3:

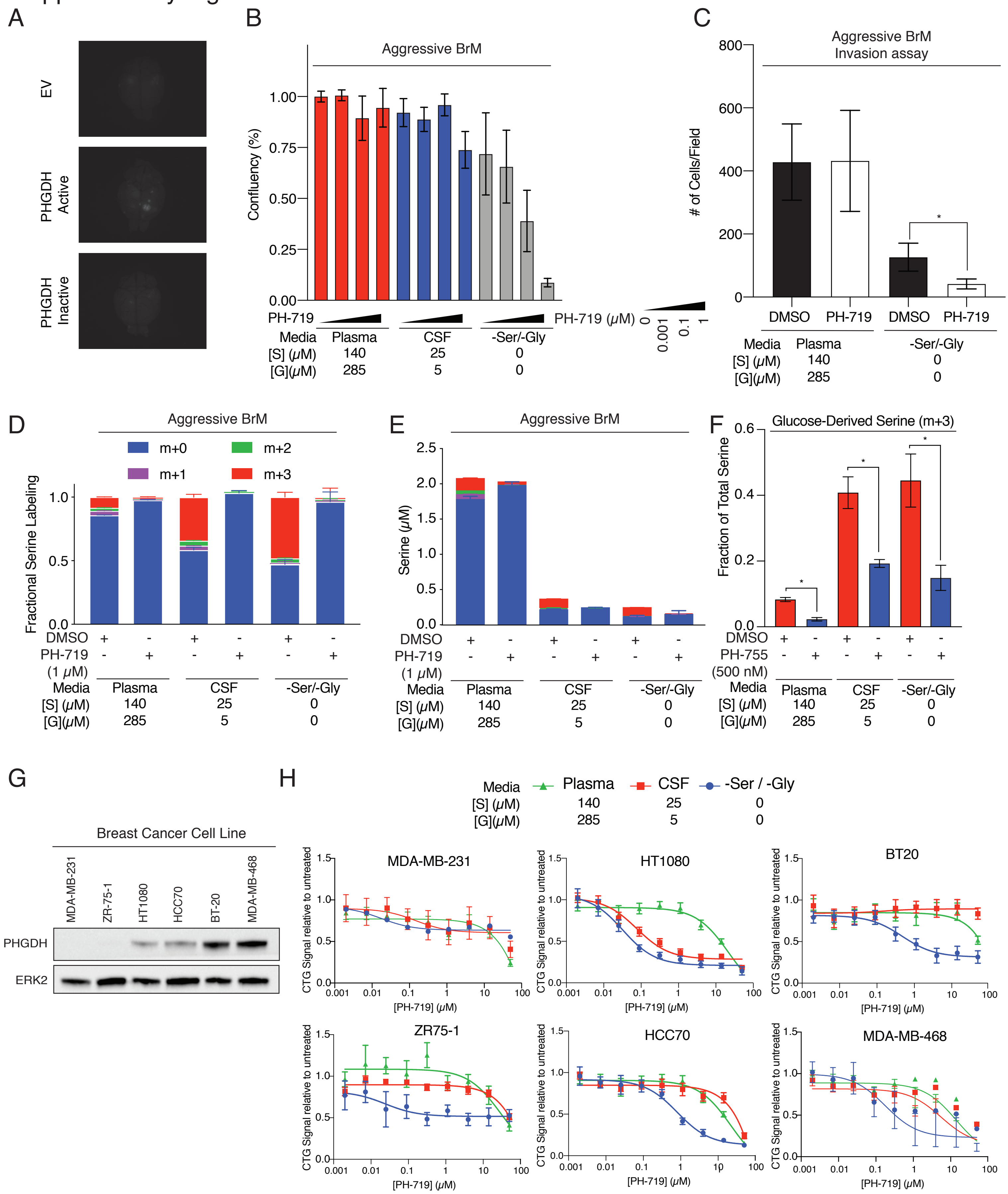

**Supplementary Figure S3, Related to Figures 3 and 4.**

**(A)** Representative images of GFP-positive brain metastatic lesions derived from tumor bearing mice injected with non-trophic MDA-MB-231 (parental) cells expressing either an empty vector (EV) control, catalytically active PHGDH, or catalytically inactive PHGDH. Active PHGDH induces brain lesion formation by parental MDA-MB-231 cells. The empty vector and inactive PHGDH fail to do so. **(B)** Proliferative capacity of aggressive brain metastatic cells cultured in media with plasma or CSF concentrations of serine and glycine, or media completely lacking serine and glycine (-Ser/-Gly), and treated with DMSO, 10nM, 100nM, or 1$\mu$M of PH-719. Decreased concentrations of extracellular serine and glycine enhance cellular responses to PHGDH inhibition. **(C)** Cell invasion assay of aggressive BrM cells cultured in media with plasma or -Ser/-Gly media, and treated with DMSO or 1$\mu$M of PH-719. Suppression of PHGDH diminishes invasive capacity through a Matrigel membrane. **(D)** Fractional labeling of U-$^{13}$C-Glucose derived serine from aggressive brain metastatic cells grown in Plasma, CSF, or -Ser/-Gly media, and treated with DMSO or 1$\mu$M of PH-719. Pharmacological inhibition of PHGDH activity abrogates cellular capacity to synthesize serine from glucose. **(E)** Serine pools, normalized to cell counts, in aggressive brain metastatic cells treated with DMSO or 1 $\mu$M PH-719. PH-719 treatment prevents the production of m+3, glucose-derived serine. **(F)** Fraction of glucose derived serine from aggressive BrM cells cultured in Plasma, CSF, or -Ser/-Gly media and treated with DMSO or 500nM of PH-755. PH-55 effectively blocks glucose derived serine synthesis. **(G)** Western blot analysis of PHGDH expression in MDA-MB-231, ZR-75-1, HT1080, HCC70, BT20, and MDA-MB-468 cells. MDA-MB-231 and ZR-75-1 cells do not express PHGDH at the protein level, but PHGDH is present in the remaining cell lines. **(H)** PH-719 dose response curves of MDA-MB-231, MDA-MB-468, HT1080, HCC70, BT20, and ZR-75-1 cells grown in plasma, CSF, or -Ser/-Gly media. Elimination of serine from the media sensitizes PHGDH-expressing cell lines to treatment with PH-719. MDA-MB-231 and ZR-75-1 cell lines, which do not express the target PHGDH, are not sensitive to treatment with PH-719 regardless of media serine and glycine concentrations.

### Supplementary Figure S4:

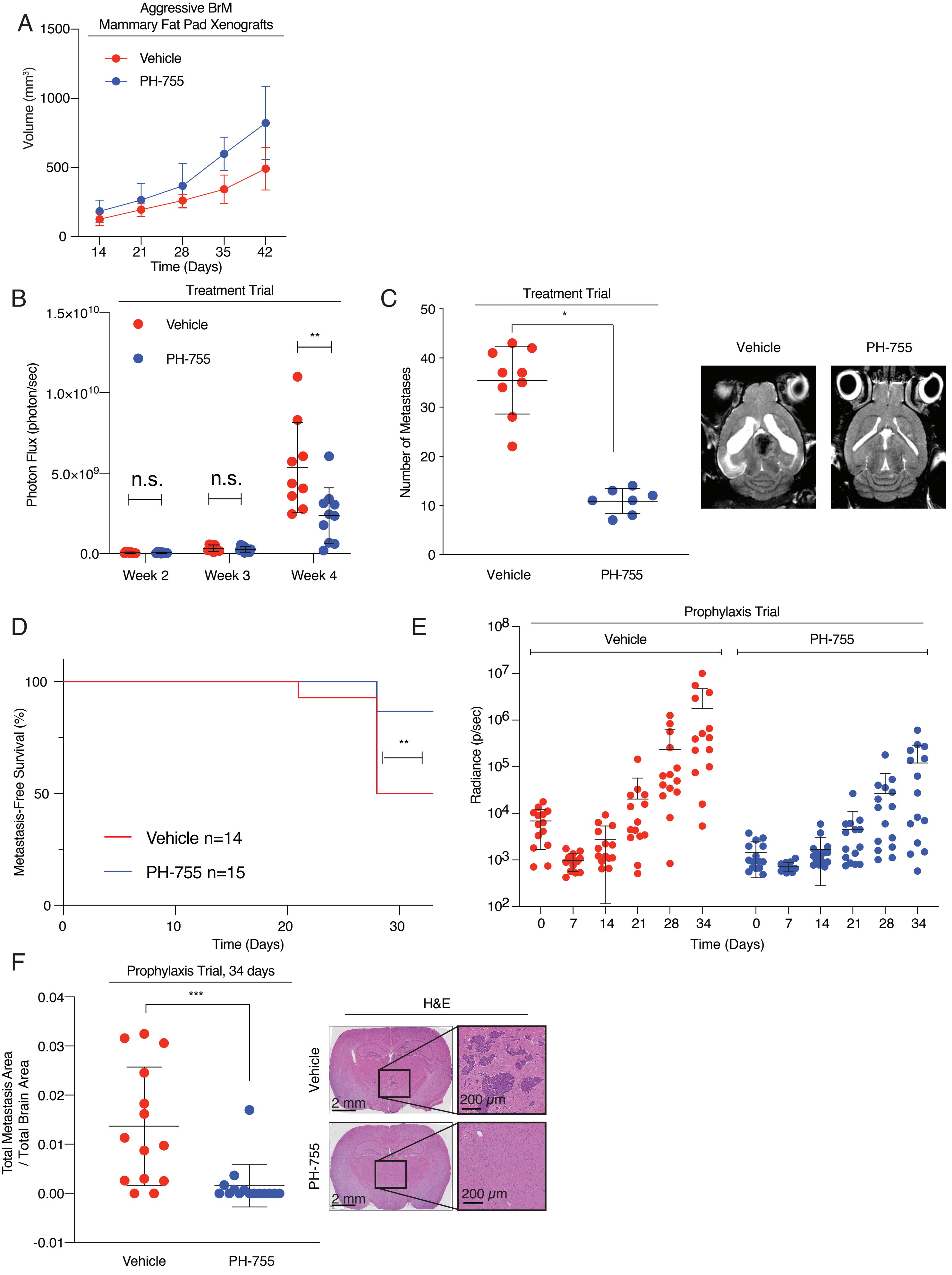

**Supplementary Figure S4, Related to Figure 4.**

**(A)** Tumor volumes of aggressive brain-metastatic cells established as xenografts in the 4th mammary fat pad (250,000 cells/injection) treated with vehicle control or 300mg/kg of PH-755 twice daily two weeks following tumor establishment. Treatment with PH-755 does not reduce mammary fat pad tumor growth.**(B)** Quantified total photon flux by bioluminescence imaging of brain metastasis bearing mice treated over time with vehicle control or 300mg/kg of PH-755 twice daily. Suppression of brain metastatic growth is observed at week 4. **(C)** Blinded quantification of brain metastatic lesions by magnetic resonance imaging (MRI) from mice treated with vehicle control or 300mg/kg of PH-755 twice daily. PH-755 suppresses the number of detected brain metastatic lesions. **(D)** Metastasis-free survival of mice from prophylaxis trial treated with vehicle or 300 mg/kg of PH-755 twice daily. Mice were defined to have brain metastases once the total brain photon flux exceeded brain photon flux measured on the day of injec-tion. Prophylactic PH-755 treatment increases metastasis-free survival at 34 days. **(E)** Quantified total photon flux by biolumi-nescence imaging of brain metastasis bearing mice from prophylaxis trial treated over 5-weeks with vehicle control or 300 mg/kg of PH-755 twice daily. PH-755 treatment reduces the brain burden of mice from day 28. **(F)** Quantification of brain metastatic lesions by H&E staining from mice in prophylaxis trial at week 5. Mice were treated with vehicle control or 300 mg/kg of PH-755 twice daily. Tumor areas were measured and quantified by two independent observers. PH-755 treatment significantly reduces the burden of brain metastases.

Supplementary Figure S5:

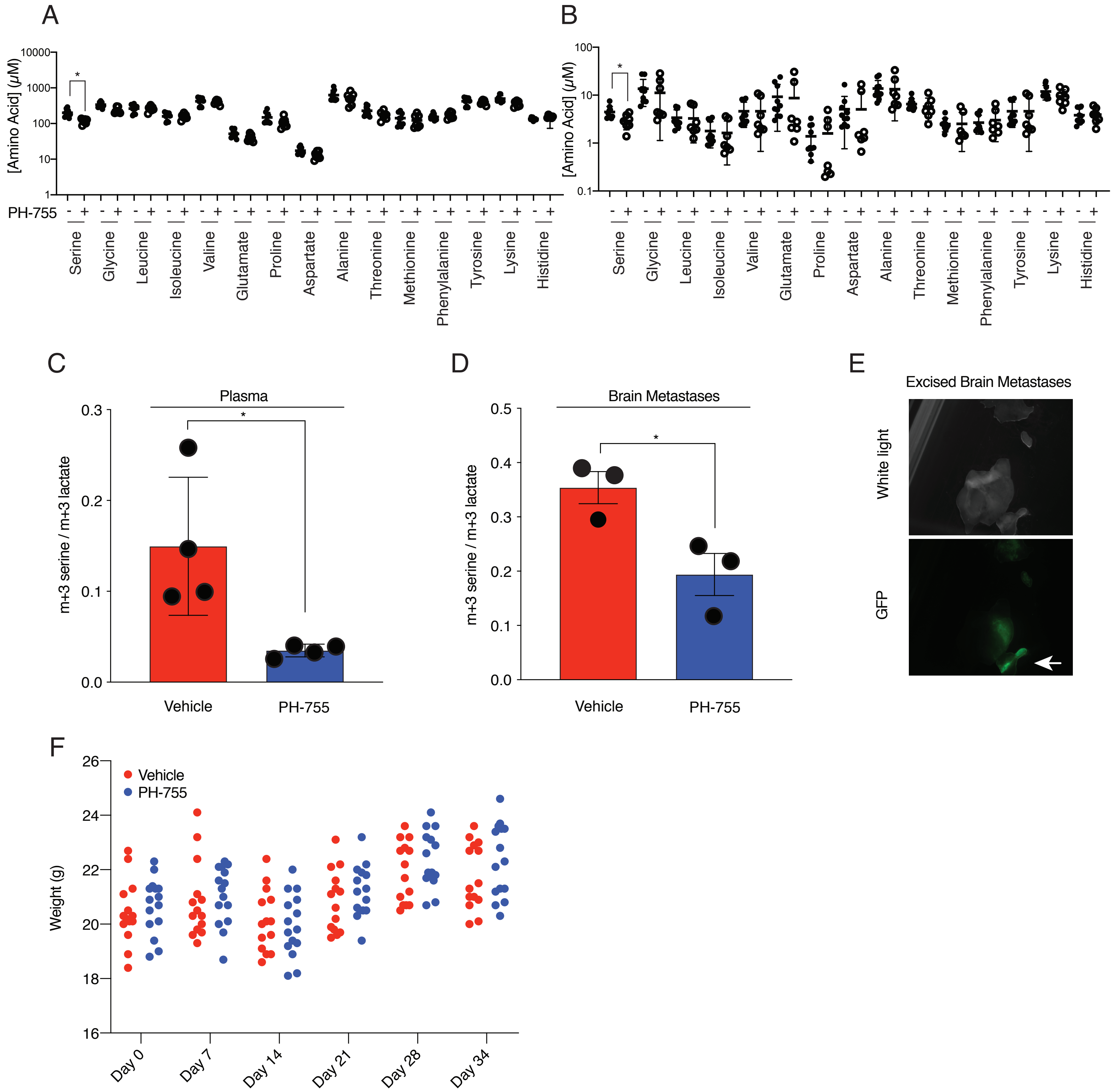

**Supplementary Figure S5, Related to Figures 4 and 5.**

**(A)** Plasma and **(B)** CSF amino acid concentrations from mice 6 hours after treatment with vehicle control or 300mg/kg of PH-755. PH-755 specifically suppresses systemic serine concentrations only. **(C)** Fraction of glucose-derived serine, normalized to fraction of glucose-derived lactate, from the plasma of mice 6 hours after treatment with vehicle control or 300mg/kg of PH-755. PH-755 effectively lowers plasma concentrations of glucose derived serine. **(D)** Fraction of glucose derived serine, normalized to fraction of glucose-derived lactate, from brain metastatic lesions isolated from mice 6 hours after treatment with vehicle control or 300mg/kg of PH-755. PH-755 lowers the fraction of glucose-derived serine in brain metastases. **(E)** Representative image of isolated GFP-positive brain metastatic lesions following excision. Arrow indicates representative tumors used for supplemental Fig. 4d, with minimal surrounding brain tissue. **(F)** Weight of individual mice plotted over the course of a 5-week prophylaxis PH-755 trial. Mice were treated with vehicle control or 300mg/kg twice daily of PH-755. PH-755 treated mice demonstrated no weight loss compared to the control.

Supplemental Table S1. shRNA sequences used in this work.

| Name | TRC Number (if applicable) | Sequence |
|---|---|---|
| shNT | N/A | CCGGCAACAAGATGAAGAGCACCAACTCGAGTTGGTGCTCTTCATCTTGTTGTTTTT |
| shGFP | N/A | CCGGACAACAGCCACAACGTCTATACTCGAGTATAGACGTTGTGGCTGTTGTTTTTTG |
| shPHGDH #1 | TRCN0000233029 | CCGGCAGGACTGTGAAGGCCTTATTCTCGAGAATAAGGCCTTCACAGTCCTGTTTTTG |
| shPHGDH #2 | TRCN0000028520 | CCGGGCTTCGATGAAGGACGGCAAACTCGAGTTTGCCGTCCTTCATCGAAGCTTTTT |
